# BDNF controls cognitive processes related to neuropsychiatric manifestations via autophagic regulation of p62 and GABA_A_ receptor trafficking

**DOI:** 10.1101/334466

**Authors:** Toshifumi Tomoda, Akiko Sumitomo, Rammohan Shukla, Yuki Hirota-Tsuyada, Hitoshi Miyachi, Hyunjung Oh, Leon French, Etienne Sibille

## Abstract

Reduced BDNF and GABAergic inhibition co-occur in neuropsychiatric diseases, including major depression. Genetic rodent studies show a causal link, suggesting the presence of biological pathways that mediate this co-occurrence. Here we show that mice with reduced Bdnf (*Bdnf*^+/-^) have upregulated expression of sequestosome-1/p62, an autophagy-associated stress response protein, and reduced surface presentation of α5 subunit-containing GABA_A_ receptor (α5-GABA_A_R) in prefrontal cortex (PFC) pyramidal neurons. Reducing *p62* gene dosage restored α5-GABA_A_R surface expression and rescued the PFC-relevant behavioral deficits of *Bdnf*^+/-^ mice, including cognitive inflexibility and sensorimotor gating deficits. Increasing p62 levels was sufficient to recreate the molecular and behavioral profiles of *Bdnf*^+/-^ mice. Finally, human postmortem corticolimbic transcriptome analysis suggested reduced autophagic activity in depression. Collectively, the data reveal that autophagy regulation through control of p62 dosage may serve as a mechanism linking reduced BDNF signaling, GABAergic deficits, and psychopathology associated with PFC functional deficits across psychiatric disorders.

**HIGHLIGHTS:** BDNF constitutively promotes autophagy in cortical pyramidal neurons

Reduced BDNF causes elevated autophagy-regulator p62 expression, leading to lower surface α5-GABA_A_R presentation

Increasing p62 levels mimics cognition-related behavioral deficits in *Bdnf*^+/-^ mice

Altered postmortem corticolimbic gene expression suggests reduced autophagic activity in depression

## INTRODUCTION

Cognitive impairment is associated with a range of psychiatric and neurological conditions (Bast et al., 2017; Knight and Baune, 2018), manifesting either as a core symptom of major mental illnesses (*e.g*., major depressive disorder [MDD], schizophrenia), as age-related decline of brain functions, or as a condition comorbid to neurodegenerative and other brain disorders. Pathobiological mechanisms underlying cognitive impairment include deficits in neural plasticity or synaptic functions (Negrón-Oyarzo et al., 2016); however, the diversity of molecular entities and multiple neurotransmitter systems implicated in synaptic function and neuroplasticity have made it difficult to pinpoint relevant central neurobiological events contributing to cognitive dysfunction, and to identify therapeutic targets that can effectively alleviate these symptoms.

Translational molecular studies have consistently reported lower expression levels of brain-derived neurotrophic factor (BDNF) in postmortem samples from subjects with MDD (Guilloux et al., 2012; Tripp et al., 2012), schizophrenia (Weickert et al., 2003; Pillai et al., 2010; Islam et al., 2017), and age-related cognitive decline (Calabrese et al., 2013; Oh et al., 2016). BDNF is a member of the neurotrophin family of growth factors (Chao, 2003) which plays a number of critical roles in the nervous system, including synaptogenesis, neurotransmission, learning, memory, and cognition (Chao et al., 2006; Lu et al., 2014). Animal models with reduced BDNF expression or activity showed disturbances of neurotransmission or neural plasticity, which likely underlie cognitive deficits observed in these models (Dincheva et al., 2012; Lu et al., 2014; Dincheva et al., 2016), consistent with a dimensional contribution of this pathway in cognitive symptoms across a range of psychiatric disorders.

BDNF primarily signals through binding to TrkB receptor and its co-receptor, p75/NTR, leading to activation or modulation of downstream signaling molecules (Chao, 2003). BDNF signaling is also regulated at the receptor level. Upon ubiquitination of TrkB and p75/NTR by TRAF6, an E3 ligase, the ubiquitin-binding adaptor protein, sequestosome-1/p62, is recruited to form a protein complex (TrkB/p75/TRAF6/Ubi/p62), which is then trafficked to an appropriate cellular compartment (*e.g*., the proteasome or lysosome for degradation, the endosome for internalization or recycling), leading to down- or up-regulation of BDNF signaling in a context-dependent manner (Sánchez-Sánchez and Arévalo, 2017). Consistent with this model, a recent study reported that TrkB is located to the autophagosome and that it can mediate retrograde transport of this organelle in neurons (Kononenko et al., 2017).

p62 is a critical adaptor protein that integrates multiple cellular processes, including growth factor signaling, ubiquitin/proteasomal system, and autophagy/lysosomal system. This occurs by interacting with signaling molecules, ubiquitinated proteins, and autophagy-related protein LC3 (Lippai and Lőw, 2014). We recently reported that p62 regulates levels of the neuronal cell surface expression of γ-aminobutyric acid (GABA)_A_ receptors (GABA_A_Rs) (Sumitomo et al., 2018a). In that study we show that in the prefrontal cortex (PFC) of mice heterozygous for *Ulk2*, an autophagy-regulatory gene, p62 protein levels are elevated as a result of attenuated autophagy, leading to sequestration of GABA_A_ receptor-associated protein (GABARAP) (Pankiv et al., 2007), an adaptor protein implicated in endocytic trafficking of GABA_A_Rs (Wang et al., 1999). This leads to selective downregulation of GABA_A_Rs on the surface of pyramidal neurons, thereby underlying cognition-related behavioral deficits observed in *Ulk2*^+/-^ mice (Sumitomo et al., 2018a).

Besides BDNF, we and others have consistently demonstrated dysfunctions of GABAergic inhibitory neurotransmission in the corticolimbic circuitry of MDD (Guilloux et al., 2012; Northoff and Sibille, 2014; Fee et al., 2017), schizophrenia (Caballero and Tseng, 2016; Hoftman et al., 2017), and age-related cognitive decline (Porges et al., 2017), suggesting a critical contribution of this inhibitory pathway to the pathophysiology of mental illnesses. Mouse–human translational studies further suggest that GABAergic changes occur downstream of reduced BDNF signaling, specifically affecting dendritic-targeting GABAergic interneurons (Guilloux et al., 2012; Tripp et al., 2012). Using human postmortem samples, we have demonstrated a positive correlation between *BDNF* and GABAergic synaptic gene expression, and in mouse models we showed that blockage of global or dendritic BDNF signaling in the PFC leads to reduction in expression of GABAergic genes mediating dendritic synaptic function (Oh et al., 2016; Oh et al., 2018), providing a causal link between reduced BDNF signaling and deregulated GABAergic neurotransmission. GABA elicits its inhibitory neurotransmission through pentameric GABA_A_Rs containing multiple subunits with diverse functional and anatomical properties, including somatically or perisomatically-targeted subunits (*e.g*., α1, α2) and dendritically-targeted subunits (*e.g*., α5).

Summing up the evidence: (1) BDNF signaling is reduced in neuropsychiatric conditions, (2) markers of GABAergic function are significantly decreased in neuropsychiatric conditions and in mice with reduced BDNF signaling, (3) dendritic BDNF transcripts and dendritically-localized α5-GABA_A_R are specifically affected in these conditions, (4) α5-GABA_A_Rs contribute to cognitive processes, and (5) autophagy-related protein (p62) regulates BDNF signaling and GABA_A_R trafficking. Accordingly, we tested the hypothesis that autophagy-related mechanisms operate downstream of BDNF, lead to the regulation of GABAergic functions preferentially through α5-GABA_A_R, and underlie deficits of cognitive processes observed across multiple neuropsychiatric conditions. We first investigated a putative new mechanism by which BDNF may control cell surface presentation of α5-GABA_A_R via regulation of p62 expression levels. We next investigated a functional and causal link between BDNF signaling and α5-GABA_A_R surface expression by bi-directional manipulation of p62 expression levels in mice and evaluation of cognitive flexibility and sensorimotor gating. We chose these two behavioral endophenotypes since they are relevant to psychiatric manifestations (Kellendonk et al., 2009; Parnaudesu et al., 2013; Swerdlow et al., 2016) and are commonly observed in BDNF mutant mice (Manning et al., 2013; Parikh et al., 2016) and α5-GABA_A_R-deficient mice (Hauser et al., 2005; Engin et al., 2013). Finally, we analyzed the relevance of the autophagy regulatory pathway to neuropsychiatric manifestations, using gene expression profiles obtained in postmortem brains of MDD and control subjects.

## RESULTS

### BDNF regulates autophagy in cortical neurons

Autophagy is a specialized membrane trafficking machinery and a major cellular recycling system primarily responsible for degrading old proteins and damaged organelles in the lysosome, which ultimately contributes to the maintenance of cellular homeostasis (Mizushima and Komatsu, 2011). Recent studies demonstrated new roles for autophagy in higher-order brain functions, through synapse pruning (Tang et al., 2014) or GABA_A_ receptor trafficking (Sumitomo et al., 2018a), suggesting that autophagy deficits represent a pathological mechanism underlying cognition-related behavioral deficits relevant to neuropsychiatric disorders, including autism (Tang et al., 2014) and schizophrenia (Sumitomo et al., 2018a, b).

As a first step to address the role of BDNF in neuronal autophagy, we prepared primary cortical neurons from transgenic mice expressing GFP-LC3, a fluorescent marker for the autophagosome (Mizushima et al., 2004), and cultured them for 16 days before treating them with BDNF (100 ng/ml) for 30 min. We observed an increase in number and size of GFP-LC3-positive punctate structures (**Figure 1A**), suggesting either *de novo* autophagosome formation due to autophagy induction, or the accumulation of LC3 due to attenuated autophagic degradation. To discriminate between these two possibilities, we performed an autophagy flux assay in primary cortical neurons (Mizushima et al., 2010) and showed that BDNF increased the autophagy flux, *i.e*., the difference between the amount of membrane-bound LC3 (*i.e*., LC3-II) seen in the presence versus the absence of lysosomal protease inhibitors, which reflects the amount of LC3 degraded through an autophagy-dependent process within the lysosome (**Figure 1B**). These results suggest that BDNF has an autophagy-inducing activity in cortical neurons.

**Figure 1.**
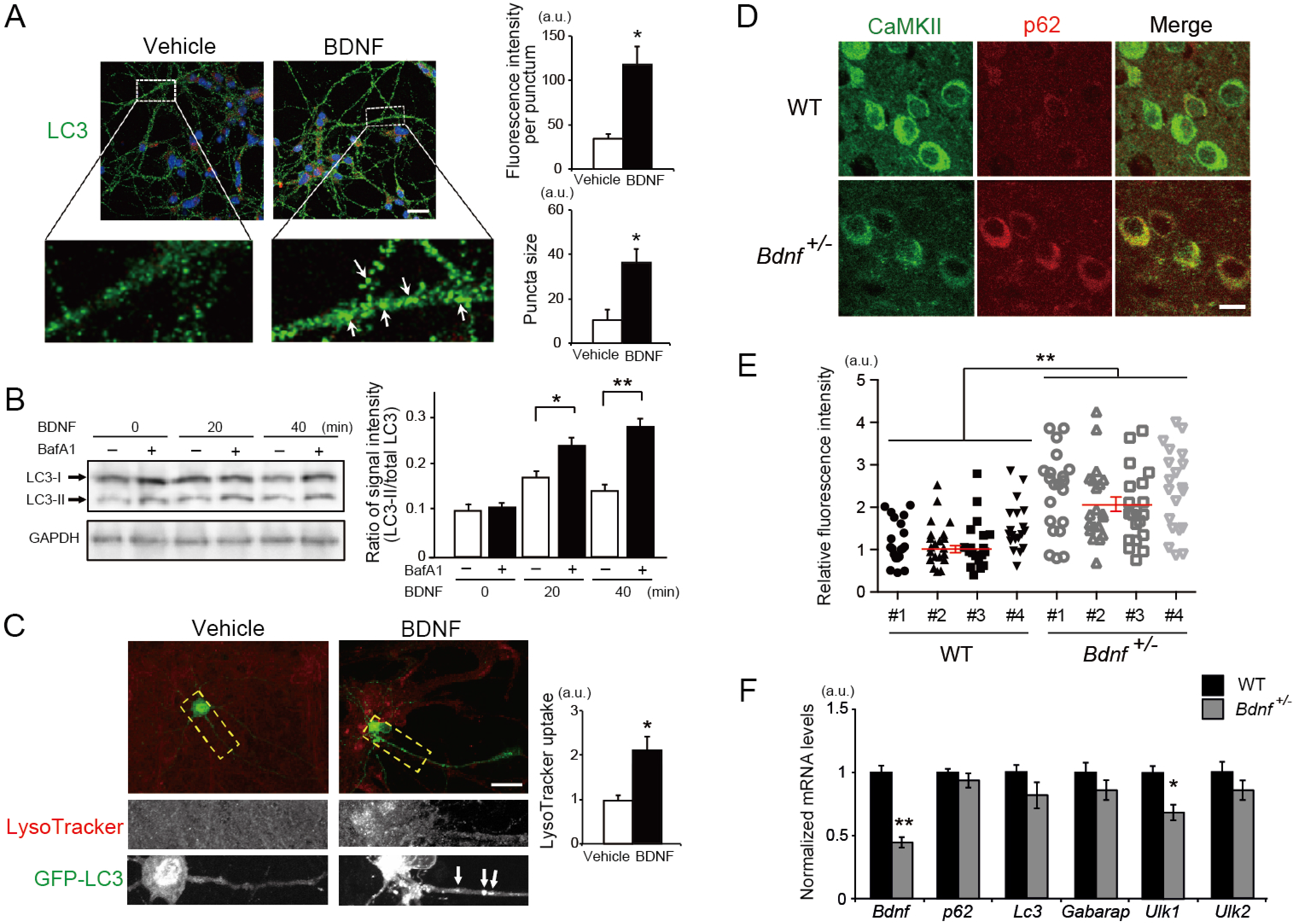
BDNF regulates autophagy and p62 protein levels in cortical pyramidal neurons. (A) Primary cortical neurons prepared from GFP-LC3 mice were treated with BDNF (100 ng/ml) or vehicle for 30 min, and the fluorescence intensities of GFP+ autophagosomes and their size were scored from >50 GFP+ puncta per condition. Scale bar, 20 μm. *p<0.05 (Kruskal-Wallis test). (B) Autophagy flux assay: primary cortical neurons were treated with BDNF (100 ng/ml) for the indicated times in the presence or absence of a lysosomal protease inhibitor (bafilomycin A1 [BafA1], 5 ng/ml) and the cell lysates were analyzed by Western blot using LC3 antibody. *p<0.05, **p<0.01 (Kruskal-Wallis test). (C) Primary cortical neurons prepared from wild-type mice were transfected with GFP-LC3 and treated with BDNF (100 ng/ml) for 30 min. LysoTracker uptake was evaluated to measure acidity of the lysosome in culture. Scale bar, 20 μm. The dotted line areas were magnified with separate LysoTracker and LC3 signals in the two lower panels. Arrows indicate GFP+ autophagosomes located in the neurite. *p<0.05 (Kruskal-Wallis test). (D, E) The PFC (layer 2/3) of *Bdnf*^+/-^ mice (*n* = 4) and their WT littermates (*n* = 4) (2 to 2.5 months of age) were immunostained with p62 and CaMKII antibodies, and p62 fluorescence intensities in CaMKII+ neurons were scored and plotted in (E). Scale bar, 20 μm. *p<0.05 (Kruskal-Wallis test). (F) Quantitative PCR analysis of autophagy-related genes expressed in the PFC of WT (*n* = 3) and *Bdnf*^+/-^ mice (*n* = 3) (2 months of age). *p<0.05, **p<0.01 (Kruskal-Wallis test).

We next tested whether BDNF could also affect the later phase of autophagy (*i.e*., maturation stage), where proteins and organelles are degraded in the acidophilic lysosome. This can be assessed by evaluating the acidity of the autophagosome/lysosomal system using LysoTracker (Mizushima et al., 2010). To facilitate simultaneous observation of autophagosome formation and maturation, primary cortical neurons prepared from wild-type (WT) mice were transfected with GFP-LC3 and then labeled with LysoTracker for the last 5 min of the culture period, 30 min after adding BDNF. BDNF markedly increased the extent of LysoTracker uptake by the soma (**Figure 1C**), indicating that BDNF promoted maturation of the autophagy/lysosomal system in neurons. Consistently, neurons with greater levels of LysoTracker uptake also exhibited increase in number and size of autophagosomes located in neurites (**Figure 1C**, arrows), suggesting a sequence of events elicited by BDNF, from autophagosome formation to maturation. Together these data show that BDNF has an autophagy-enhancing activity in cultured cortical neurons.

### Reduced BDNF expression leads to elevated p62 levels in cortical pyramidal neurons

To address the endogenous activity of BDNF in the regulation of autophagy *in vivo*, we quantitated the level of p62 protein in the medial prefrontal cortex (mPFC) of mice with reduced *Bdnf* levels (*Bdnf*^+/-^ mice). In this brain region, BDNF is predominantly produced by CaMKII-positive pyramidal neurons and functions as an autocrine and paracrine factor to modulate the activity of neighboring excitatory and inhibitory neurons (Gorba and Wahle, 1999). p62 is used as an *in vivo* marker of autophagic activity, because it is the obligatory adaptor protein selectively targeted for autophagic degradation (Mizushima et al., 2010). The results show a significant increase in p62 protein levels in CaMKII-positive neurons of *Bdnf*^+/-^ mice, as compared to WT control mice (**Figure 1D** and **1E**). This increase in protein level was not paralleled by an increase in transcriptional activity of the *p62* gene (**Figure 1F**), implying that the increased levels of p62 may result from reduced rates of protein degradation. These results suggest decreased autophagic activity in the presence of reduced BDNF levels, consistent with the hypothesis that BDNF constitutively enhances autophagy. To obtain further evidence in support of reduced autophagy in *Bdnf*^+/-^ mice, we quantitated levels of expression of additional genes in this pathway. Expression of several autophagy regulatory genes (*e.g., Lc3, Gabarap, Ulk2*) remained unchanged, whereas expression of *Ulk1*, a gene critical to autophagy induction (Mizushima and Komatsu, 2011), was significantly downregulated by ~25% (**Figure 1F**). Together the data demonstrate a constitutive role for BDNF in enhancing autophagy, manifested by attenuated autophagy and persistent upregulation of p62 protein expression in the PFC of *Bdnf*^+/-^ mice.

### Elevated p62 expression in *Bdnf*^+/-^ cortical neurons causes downregulated surface presentation of α5-GABA_A_R

We recently reported that, similar to *Bdnf*^+/-^ mice, p62 protein expression levels are elevated in pyramidal neurons of the PFC in genetically-engineered mice with attenuated autophagy (*Ulk2*^+/-^), leading to downregulation of neuronal surface expression of GABA_A_ receptors through sequestration of GABARAP, an adaptor protein responsible for endocytic trafficking of GABA_A_ receptors (Sumitomo et al., 2018a). Besides, we previously showed that reduced BDNF signaling in the PFC causes decreased expression of GABA synaptic genes, most notably α5-GABA_A_R (Oh et al., 2016), a GABA_A_ receptor subtype specifically expressed in the dendrites of pyramidal neurons (Fritschy and Mohler, 1995). These results suggest that BDNF affects GABA neurotransmission through regulation of α5-GABA_A_R levels in the dendrites of pyramidal neurons, consistent with *Bdnf*^+/-^ mice exhibiting reduced amplitude and frequency of inhibitory miniature currents (mIPSC) in cortical and thalamic neurons (Laudes et al., 2012). We therefore reasoned that elevated p62 protein levels in *Bdnf*^+/-^ neurons may influence the surface presentation of α5-GABA_A_R.

To test this hypothesis, we performed surface biotinylation of cultured cortical neurons and showed that cell surface α5-GABA_A_R protein levels were reduced in *Bdnf*^+/-^ neurons, as compared with WT control neurons, with no significant changes in total levels of α5-GABA_A_R, or in both total and surface levels of the glutamate receptor NR1 (**Figure 2A**). Notably, reducing the *p62* gene dosage in *Bdnf*^+/-^ neurons, using cortical neurons from *Bdnf*^+/-^;*p62*^+/-^ mice, restored α5-GABA_A_R surface levels to WT levels (**Figure 2A**), suggesting that p62 is a critical adaptor mediating BDNF-induced altered surface presentation (*i.e*., trafficking) of α5-GABA_A_R.

**Figure 2.**
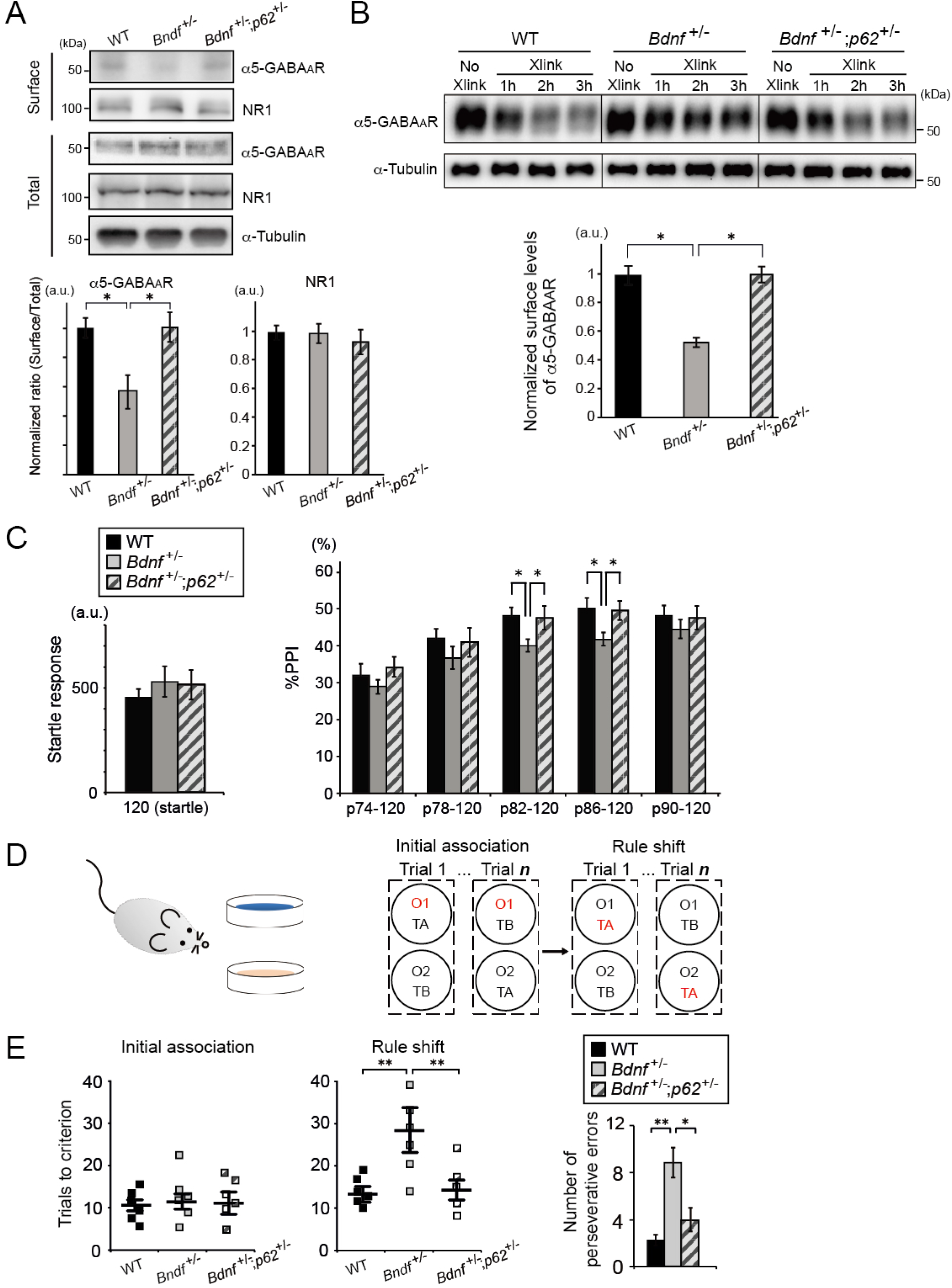
Decreased surface presentation of α5-GABA_A_R and behavioral deficits in *Bdnf*^+/-^ mice are rescued by reducing *p62* gene dosage. (A) Surface biotinylation of primary cortical neurons prepared from WT, *Bdnf*^+/-^, or *Bdnf*^+/-^;*p62*^+/-^ mice, analyzed by Western blot using the indicated antibodies. Surface levels of expression were normalized by the total levels of expression for each genotype. Surface levels of α5-GABA_A_R were significantly reduced in *Bdnf*^+/-^ neurons compared to the other genotypes, whereas those of NR1 subunit of glutamate receptors were equivalent across the three genotypes. The assays were performed in triplicate. *p<0.05 (Kruskal-Wallis test). (B) BS3 cross-linking assays using the PFC extracts from WT, *Bdnf*^+/-^, or *Bdnf*^+/-^;*p62*^+/-^ mice. Levels of non-crosslinked α5-GABA_A_R (~50 kDa) were normalized to levels of α-Tubulin at each time point. Differences in the levels of non-crosslinked α5-GABA_A_R at a given time point versus those of the control sample (No Xlink: no cross-linker added) represent the amount of surface α5-GABA_A_R that underwent mobility shift toward a higher molecular weight range due to covalent cross-linking with anonymous cell surface proteins. Cross-linking reaction reached plateau in 2–3 h in our assay conditions. Calculated surface α5-GABA_A_R levels (= levels at No Xlink – levels at 3 h Xlink) in *Bdnf*^+/-^ mice were significantly reduced compared to the other genotypes. *p<0.05 (Kruskal-Wallis test). See also **Figure S1** for cross-linking assays with α1- and α2-GABA_A_Rs. (C) The amplitude of startle response and the percentage of PPI were evaluated for WT (*n* = 10), *Bdnf*^+/-^ (*n* = 10), and *Bdnf*^+/-^;*p62*^+/-^ mice (*n* = 10). No significant difference in startle response (left panel; *F*_2,27_ = 0.378, *P* = 0.9243, one-way ANOVA). PPI was reduced in *Bdnf*^+/-^ mice compared to WT mice, and rescued to control levels in *Bdnf*^+/-^;*p62*^+/-^ mice for the prepulse–pulse pair of trials (p82–120 and p86–120 dB) (right panel; *F*_2,27_ = 9.775, *P* < 0.001, two-way ANOVA with repeated measures; *p<0.05, Bonferroni post-hoc test). (D) Schematic diagram of the rule shift assay: Mice were habituated to food, feeding apparatus, different odor cues (O1, O2, etc.; *e.g*., coriander versus garlic powder) and texture cues (TA, TB, etc.; *e.g*., fine versus coarse digging media) prior to testing, and then food-deprived a day before the assays. Mice were initially trained in a sequence of trials to associate a food reward with a specific stimulus (*i.e*., either an odor or a digging medium; a stimulus associated with food reward is shown in red). A varying combination of stimulus and food reward was presented to mice per trial. Eight consecutive correct responses to the food reward were considered reaching criterion (*i.e*., successful establishment of association between the stimulus and the food reward), and the number of trials to reach criterion were scored for each mouse tested, before and after rule shifting (*e.g*., from an odor cue to a different texture cue to predict reward). (E) Numbers of trials to criterion were scored for WT (*n* = 6), *Bdnf*^+/-^ (*n* = 6), and *Bdnf*^+/-^;*p62*^+/-^ mice (*n* = 6) during the initial association phase, as well as the rule shift phase of the assays. No significant difference during the initial association phase (*F*_2,15_ = 1.25, *P* = 0.934). Statistical significance during the rule shift phase: *F*_2,15_ = 9.93, *P* < 0.001 (one-way ANOVA); \*\**p*<0.01 (Bonferroni post-hoc test). During the rule shift phase, *Bdnf*^+/-^ mice made a greater number of perseverative errors than the other genotypes. *p<0.05, **p<0.01 (Kruskal-Wallis test).

To validate this finding *in vivo*, we performed receptor cross-linking assays using bis(sulfosuccinimidyl)suberate (BS3), a membrane-impermeable chemical cross-linker (Boudreau et al., 2012). As the cross-linking reaction proceeds in the presence of BS3, only the fraction of receptors expressed on the plasma membrane surface are expected to be covalently cross-linked with anonymous cell surface proteins, thereby transforming into higher molecular weight species, while the rest of the receptors associated with the endomembrane would remain intact, maintaining their original molecular weights. The PFC from WT and *Bdnf*^+/-^ mice were subjected to the cross-linking reaction and the levels of a series of GABA_A_R subunits (*i.e*., α1, α2, and α5) were measured as a function of time (1–3 h). The results showed equivalent surface levels of α1- or α2-GABA_A_R (**Figure S1**), and a significant time-dependent decrease in the levels of intact α5-GABA_A_R (~50 kDa) in Bdnf^+/-^ mice; specifically ~53% of total α5-GABA_A_R were estimated to be expressed on the cell surface in WT PFC (**Figure 2B**, left), whereas a lesser extent (~25%) of α5-GABA_A_R were presented on the cell surface under conditions of reduced *Bdnf*^+/-^ levels (**Figure 2B**, middle). In contrast, ~56% of α5-GABA_A_R were observed in the PFC of *Bdnf*^+/-^ in which p62 levels were genetically reduced (*Bdnf*^+/-^;*p62*^+/-^) (**Figure 2B**, right), demonstrating that reducing the *p62* gene dosage restored the surface expression of α5-GABA_A_R to a level equivalent to WT mice. Collectively, the data demonstrate that decreased BDNF expression results in specific reduction in surface presentation of α5-GABA_A_R through elevated p62 expression in the PFC.

### Elevated p62 expression in *Bdnf*^+/-^ mice mediates behavioral deficits relevant to PFC dysfunction

We then investigated the potential role of elevated p62 expression in PFC-relevant brain functions of *Bdnf*^+/-^ mice, such as information processing and cognition. Previous studies reported that *Bdnf*^+/^ mice have reduced prepulse inhibition (PPI) of acoustic startle response (Manning et al., 2013), demonstrating a role of BDNF in sensorimotor gating function. This mechanism of filtering sensory information to render appropriate motor responses has been shown to rely in part on the function of the cortical circuitry involving mPFC (Swerdlow et al., 2016). In agreement with the previous study, we first confirmed normal startle response and reduced PPI levels in *Bdnf*^+/-^ mice (**Figure 2C**). We next showed that reducing the *p62* gene dosage in *Bdnf*^+/-^ mice (*i.e*., using *Bdnf*^+/-^;*p62*^+/-^ mice) rescued the PPI deficits back to levels observed in control WT mice (**Figure 2C**). Together these results suggest that decreased expression of BDNF leads to sensorimotor gating deficits through elevated p62 expression.

*Bdnf*^+/-^ mice were recently shown to exhibit reduced cognitive flexibility in a visual discrimination task (Parikh et al., 2016). To further address deficits in cognitive flexibility in *Bdnf*^+/-^ mice, we used a rule shifting paradigm (Bissonette et al., 2008; Cho et al., 2015), in which mice were initially trained to associate food reward with a specific stimulus (*i.e*., either an odor or a digging medium) and subsequently evaluated for cognitive flexibility by changing the type of stimulus that predicts the reward (**Figure 2D**). *Bdnf*^+/-^ and WT mice learned the association rule in a similar number of trials during the initial association phase of trials; however, *Bdnf*^+/-^ mice required significantly higher numbers of trials to shift their behavior during the rule shifting phase of trials (**Figure 2E**). When the *p62* gene dosage was reduced in *Bdnf*^+/-^ mice (*i.e*., in *Bdnf*^+/-^;*p62*^+/-^), the mice showed cognitive performance indistinguishable from that of controls (**Figure 2E**), together suggesting that decreased expression of BDNF results in cognitive deficits through elevated p62 expression.

### Elevated p62 expression is sufficient to cause downregulation of surface α5-GABA_A_R expression and behavioral deficits

To further address the causal role of elevated p62 expression in the regulation of α5-GABA_A_R surface expression and the associated behavioral changes, we generated p62-transgenic (Tg) mice, in which p62 transgene expression was driven by the CaMKII promoter. Among three Tg lines established, two lines (#1, #3) showed ~60% increase in p62 protein expression in the PFC, while one line (#2) failed to overexpress p62, as evaluated by Western blot (**Figure 3A**). BS3 cross-linking assays demonstrated reduced levels of surface α5-GABA_A_R expression in the PFC of the overexpressing p62-Tg line (#1) compared to WT, whereas the non-overexpressing line #2 displayed surface α5-GABA_A_R expression equivalent to WT levels (**Figure 3B**).

**Figure 3.**
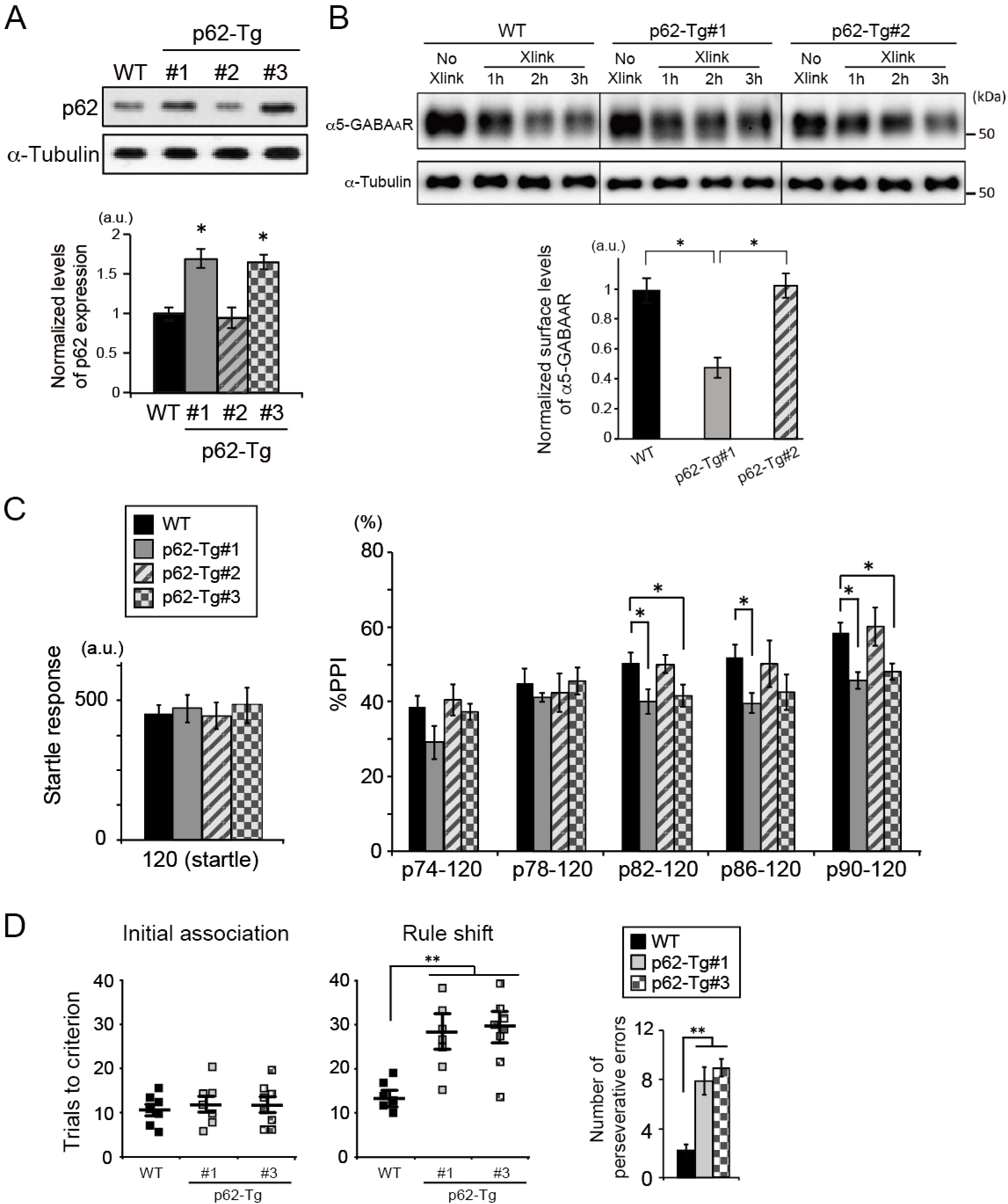
Elevated p62 expression is sufficient to cause downregulation of surface α5-GABA_A_R expression and behavioral deficits. (A) Generation of CaMKII-p62-transgenic mouse lines. Among 3 lines established, 2 lines (#1, #3) showed ~60% higher levels of p62 protein expression in the PFC when compared with WT, whereas the line #2 showed no apparent increase in p62 expression. *p<0.05 (Kruskal-Wallis test). (B) BS3 cross-linking assays using the PFC extracts from WT, p62-Tg#1 and Tg#2 mice. Levels of non-crosslinked α5-GABA_A_R (~50 kDa) were normalized by levels of α-Tubulin at each time point. Calculated surface α5-GABA_A_R levels (= levels at No Xlink – levels at 3 h Xlink) in p62-Tg#1 mice were significantly reduced compared to the other genotypes. *p<0.05 (Kruskal-Wallis test). (C) The amplitude of startle response and the percentage of PPI were evaluated for WT (*n* = 10), p62-Tg#1 (*n* = 10), p62-Tg#2 (*n* = 10), and p62-Tg#3 mice (*n* = 10). No significant difference in startle response (*F*_3,36_ = 0.726, *P* = 0.7121, one-way ANOVA). Statistical significance for %PPI; *F*_3, 36_ = 12.474, *P* < 0.001 (two-way ANOVA with repeated measures); *p<0.05 (Bonferroni post-hoc test). (D) Numbers of trials to criterion were scored for WT (*n* = 7), p62-Tg#1 (*n* = 7), and p62-Tg#3 mice (*n* = 8) during the initial association phase, as well as the rule shift phase of the assays. No significant difference during the initial association phase (*F*_2,19_ = 1.968, *P* = 0.936). Statistical significance during the rule shift phase: *F*_2,19_ = 12.56, *P* < 0.001 (one-way ANOVA); **p< 0.01 (Bonferroni post-hoc test). During the rule shift phase, p62-Tg mice made a greater number of perseverative errors than WT. **p<0.01 (Kruskal-Wallis test).

We next evaluated sensorimotor gating in the p62-Tg lines. p62-Tg lines #1 and 3 exhibited reduced levels of PPI, whereas the non-overexpressing line #2 had PPI levels that were not different from WT levels (**Figure 3C**). Similarly, in the cognitive flexibility test, mice from the overexpressing p62-Tg line (#1, #3) learned the association rule during the initial association phase of trials in a similar manner as WT, but were impaired during the rule shifting phase of trials (**Figure 3D**).

Together these data demonstrated that elevated p62 levels in CaMKII-expressing pyramidal neurons in the PFC and corticolimbic areas are sufficient to replicate the molecular and behavioral phenotypes of *Bdnf*^+/-^ mice; specifically, reduced surface expression of α5-GABA_A_R and PFC-relevant behavioral deficits.

### Human postmortem gene expression profiles suggests altered autophagy in depression

Although reduced levels of expression and/or functioning of BDNF and α5-GABA_A_R are consistently reported in the brains of several neuropsychiatric conditions, including MDD (Guilloux et al., 2012; Fee et al., 2017), the cellular machinery that may underlie these changes remains to be elucidated. On the basis of our recent reports showing attenuated neuronal autophagy in several mouse models for neuropsychiatric disorders such as schizophrenia (Sumitomo et al., 2018a, b), as well as the present data demonstrating elevated p62 levels in *Bdnf*^+/-^ mouse model, we hypothesized that alteration in autophagy machinery may contribute to cellular deficits present in MDD.

To test this hypothesis, we analyzed genome-wide differential expression statistics from a meta-analysis of eight large-scale expression datasets in corticolimbic areas of 51 MDD patients and 50 controls (Ding et al., 2015). Autophagy-related genes were defined by gene ontology (GO) (Huang et al., 2009) and combined into “autophagy-enhancing” and “autophagy-attenuating” gene lists (See details in **Tables S1** and **S2**). Of the 95 genes included in the “autophagy-enhancing” gene list, the expression of one gene was significantly increased (p<0.05), and 8 were significantly decreased (p<0.05) in MDD compared to controls (**Table S1**). By contrast, of the 38 genes included in the “autophagy-attenuating” gene list, the expression of 3 genes showed a significant increase and 2 genes a significant decrease in MDD compared to the control cohorts (**Table S2**). At the group level, an area under the curve (AUC) analysis revealed a significant difference (p<0.01) (**Figure 4A**), namely between an over-representation of upregulated autophagy-attenuating genes (AUC=0.60; 0.5 meaning no change) and a slight non-significant under-representation of downregulated autophagy-enhancing genes (AUC=0.44) in the MDD brain (**Figure 4B**). An alternate analysis using ranking of gene changes (gene set enrichment analysis, GSEA) of the same data showed no significant enrichment for autophagy-attenuating genes (**Figure 4C**, red line), but a significant enrichment in downregulation for autophagy-enhancing genes compared to controls (**Figure 4C**, green line, p=0.008). Together these results suggest reduced autophagy at the transcriptome level in MDD.

**Figure 4.**
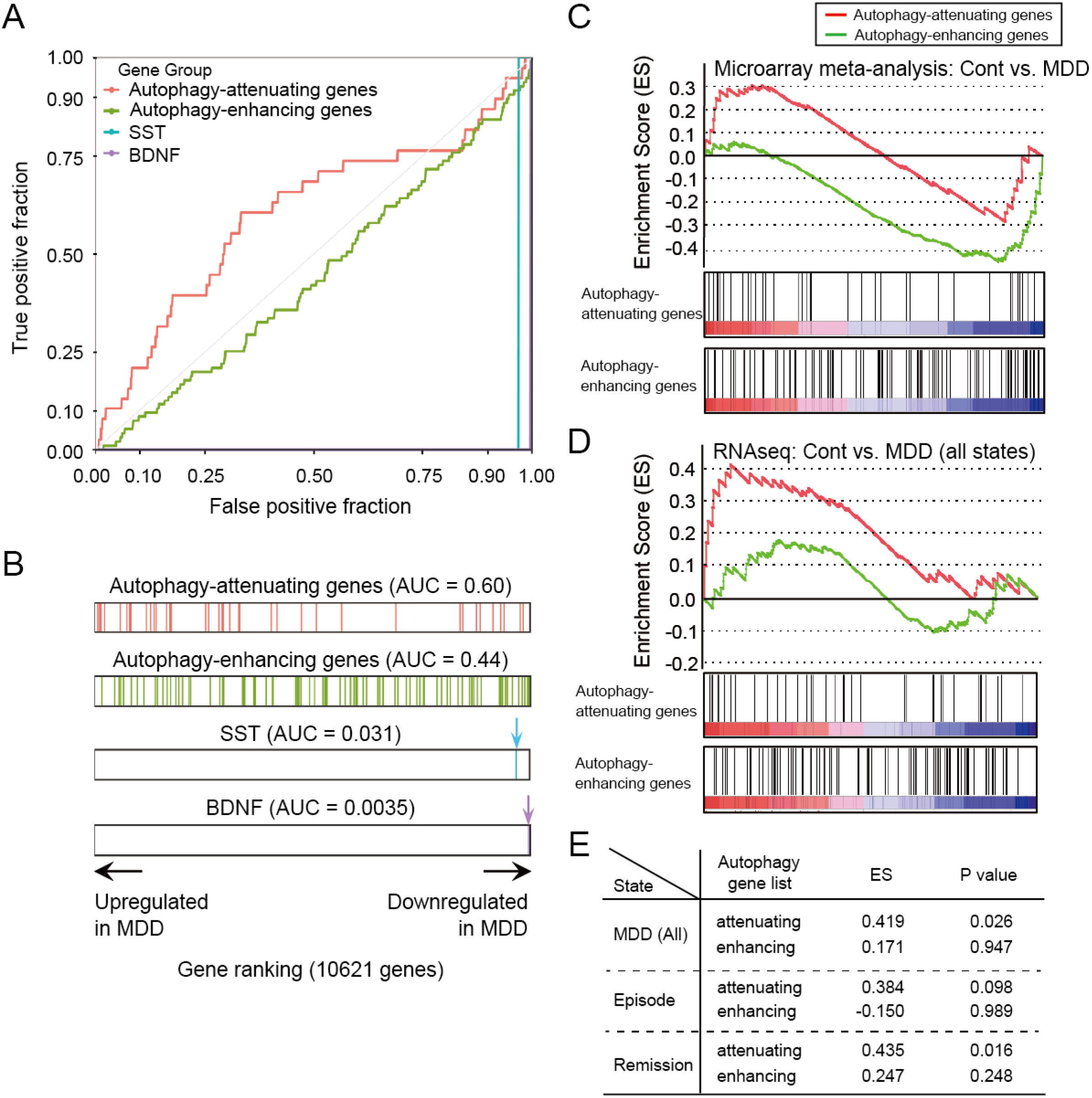
Gene expression profiles in human postmortem brains suggest reduced autophagy in MDD. (A) Expression levels of the autophagy-enhancing (95 genes, **Table S1**) versus the autophagy-attenuating (38 genes, **Table S2**) gene sets in corticolimbic areas of MDD postmortem brain samples were compared with those of the control cohorts. Overall expression levels of the autophagy-attenuating genes in MDD, as represented by the area under the curve (AUC = 0.60) were significantly different from those of the autophagy-enhancing genes (AUC = 0.44) (p<0.01). (B) The autophagy-attenuating genes in MDD clustered toward upregulated, whereas the autophagy-enhancing genes in MDD clustered toward downregulated expression. As internal reference, the ranking of the *BDNF* and *Somatostatin* (*SST*) genes were plotted, both of which have been reported to be significantly downregulated in corticolimbic areas of MDD subjects (Ding et al., 2015), as represented by the AUC values of 0.0035 and 0.031, respectively. (C) GSEA on gene expression profiles in MDD versus control subjects obtained from meta-analysis of microarray datasets shows significant enrichment in downregulation for autophagy-enhancing genes (green line) (p=0.008) but no significant enrichment for autophagy-attenuating genes (red line). (D) GSEA on gene expression profiles in MDD (all episode and remission states combined) versus control subjects obtained via RNAseq shows a significant enrichment in upregulation for autophagy-attenuating genes (red line) (p=0.026) but no significant enrichment for autophagy-enhancing genes (green line). See **Figure S2** for MDD cohorts used in RNAseq. (E) Summary of GSEA results on expression profiles of the autophagy-enhancing and autophagy-attenuating gene sets in all or disease state-specific MDD obtained via RNAseq as shown in **Figure 4D, Figures S3** and **S4**. The autophagy-attenuating gene set shows a significant enrichment in all combined MDD and in remitted cohorts or the trend level of enrichment during episodes, whereas the autophagy-enhancing gene set shows no significant enrichment regardless of disease states.

To further investigate autophagy-related gene expression profiles in different cohorts of MDD subjects, we performed RNAseq in the subgenual anterior cingulate cortex (sgACC) (Brodmann area 25) of postmortem samples obtained from four cohorts at different states of MDD (1: single episode, 2: first remission, 3: recurrent episode, and 4: second remission) and one control cohort (n=15–20/group; **Figure S2**) as described (Scifo et al., 2018). Applying GSEA to the data replicated a significant enrichment in upregulation for autophagy-attenuating genes in the combined MDD cohorts regardless of episode/remission status (1+2+3+4) compared to controls (**Figure 4D**, red line; **Figure 4E**, p=0.026). This enrichment in upregulation for the autophagy-attenuating gene set was also significant in remitted MDD subjects (cohorts 2+4) (**Figure S3**, red line; **Figure 4E**, p=0.016), but only at the trend level in currently-depressed subjects (**Figure S4**, red line; **Figure 4E**, p=0.098), suggesting either state-specific changes or reduced analytical power in smaller cohorts. No enrichment was observed for the autophagy-enhancing gene set in the combined or separate cohorts (**Figure 4D, Figures S3** and **S4**, green line; **Figure 4E**).

Collectively, the GSEA results on the distinct cohorts and platforms (microarray and RNAseq datasets) are in good overall agreement and suggest persistent autophagy attenuation in MDD.

## DISCUSSION

Building on our previous study showing a causal link between reduced BDNF signaling and selective attenuation of GABAergic gene expression in psychiatric disorders, such as MDD (Guilloux et al., 2012; Tripp et al., 2012; Oh et al., 2018) or during aging (Oh et al., 2016), we now demonstrate a novel mechanism by which (1) BDNF regulates autophagy in PFC pyramidal neurons and (2) reduced BDNF signaling negatively impacts GABA functions via autophagy-related control of GABA_A_ receptor trafficking, and show that (3) these changes underlie behavioral manifestations that are relevant to PFC-mediated symptoms of psychiatric disorders. These results show for the first time that control of p62 levels, a molecule implicated in autophagic control of cellular function, is a key molecular event linking BDNF signaling, GABAergic neurotransmission, and specific behavioral manifestations related to PFC functions. Collectively, the results suggest the presence of coordinated biological processes linking BDNF to the maintenance of neuroplasticity and excitation–inhibition balance in the PFC. As we further show that gene implicated in attenuating autophagy tend to be upregulated in MDD, reduced autophagy may contribute to the pathology of the illness and mediate reduced GABAergic function downstream of reduced BDNF signaling.

The current study identifies p62, an autophagy regulator of cellular homeostatic processes, as a link between BDNF signaling and GABAergic function. p62 was originally identified as a protein induced by cellular stress conditions, and subsequently shown to function as an adaptor protein that integrates multiple cellular processes, including the autophagy/lysosomal pathway (Lippai and Lőw, 2014). Autophagy is activated in response to a range of cellular stresses (*e.g*., depletion of nutrients and energy, misfolded protein accumulation, oxidative stress) and mitigates such stresses to maintain cellular homeostasis (Mizushima and Komatsu, 2011). Using genetic mouse and cell models, the present study, as well as our recent report (Sumitomo et al., 2018a), demonstrated that autophagy can also control cell-to-cell signaling through regulation of surface expression of receptors mediating chemical inhibition. Thus, autophagy has the potential to serve as a homeostatic cellular machinery in the context of biological disturbances associated with brain disorders. Indeed, the gene expression data obtained from the brains of human subjects suggests that the overall activity of the autophagy machinery is attenuated in MDD, and the genetic rodent model studies suggest that elevated p62 expression may partly mediate the link between reduced BDNF signaling and GABAergic disruption through reduced GABA_A_ receptor trafficking.

Reduced BDNF expression and deregulated GABA transmission frequently co-occur with psychiatric disorders, including MDD (Guilloux et al., 2012) and schizophrenia (Lewis et al., 2005), and during normal aging (Oh et al., 2016), suggesting a shared biological mechanism across these conditions. We previously demonstrated that reduced activity of BDNF, predominantly produced by pyramidal neurons, leads to reduced expression levels of presynaptic genes (*e.g., Gad1, SLC32A1*) and of neuropeptide genes (*e.g., SST, neuropeptide Y, cortistatin*) expressed in neighboring GABAergic inhibitory neurons targeting pyramidal neuron dendrites (Oh et al., 2016), together suggesting a paracrine mode of BDNF action responsible for attenuated GABA signaling (**Figure 5**, right). Moreover, expression levels of *Gabra5*, a gene encoding α5-GABA_A_R that is predominantly localized to the dendritic compartment of pyramidal neurons, were among the most significantly downregulated, suggesting an autocrine mode of BDNF action contributing to reduced expression of GABA-related genes. Both modes of transcriptional mechanisms, coupled with the attenuated GABA_A_ receptor trafficking through the autophagy regulator, as demonstrated in the current study (**Figure 5**, left), are expected to synergistically reduce GABA neurotransmission across pre- and post-synaptic compartments.

**Figure 5.**
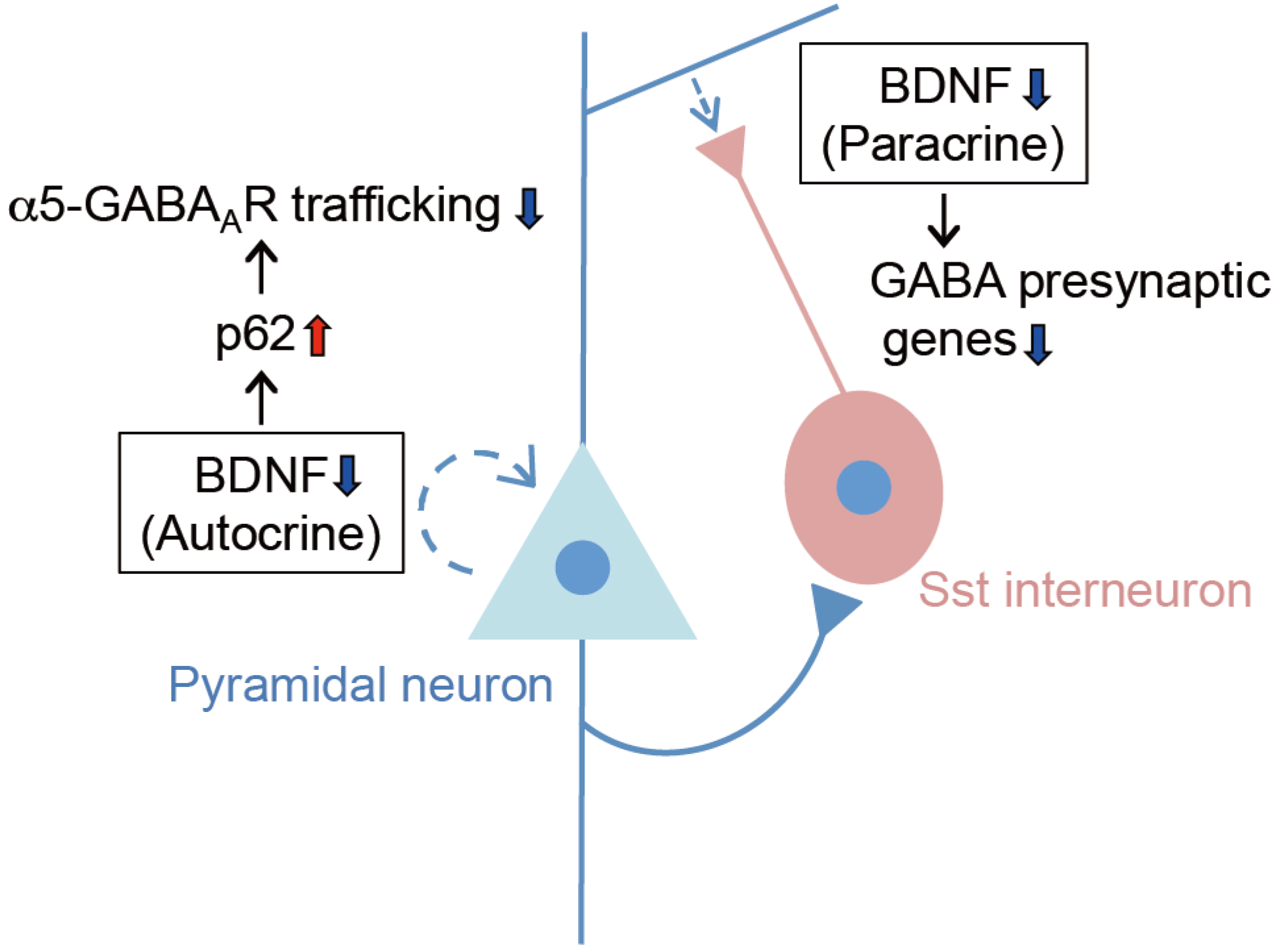
Two modes of mechanisms underlying GABAergic dysfunction following reduced BDNF signaling. Reduced BDNF expression or signaling in cortical pyramidal neurons, due to chronic stress or other neuropsychiatric insults, leads to GABA dysfunction through transcriptional suppression of GABA synapse genes in neighboring inhibitory neurons (paracrine mode) (Oh et al., 2016; Oh et al., 2018) and also via reduced surface presentation of α5-GABA_A_R in pyramidal neurons (autocrine mode), as demonstrated in this study, together contributing to cognitive and other behavioral deficits relevant to neuropsychiatric disorders, including MDD.

The functional link between reduced BDNF signaling and GABA deficits across psychiatric disorders also suggests it may contribute to common behavioral endophenotypes across disorders. Cognitive function is primarily controlled via corticolimbic mechanisms, in part through regulation of excitatory and inhibitory neurotransmission (Kellendonk et al., 2009; Caballero and Tseng, 2016), and a number of synaptic modulators have been implicated in these processes, including BDNF and α5-GABA_A_R. Notably, reduced expression or activity of BDNF or α5-GABA_A_R in mice commonly manifests as cognitive inflexibility and sensorimotor gating deficits (Hauser et al., 2005; Engin et al., 2013; Manning and van den Buuse, 2013; Parikh et al., 2016). This suggests these molecules represent a network of regulators underlying cognitive processes commonly affected across psychiatric conditions.

There are several limitations to this study. Given the multiple roles and binding partners of p62 adaptor protein, it is unlikely that α5-GABA_A_R is the only receptor system or cellular target affected by elevated p62 levels, and additional cellular machineries may further contribute to the behavioral deficits in *Bdnf*^+/-^ mice. Nonetheless we found here and in our prior transcriptomic studies that α5-GABA_A_Rs are preferentially affected, as opposed to α1-GABA_A_Rs or α2-GABA_A_Rs for instance, which are localized to somatic and perisomatic cellular compartments. Based on the predominant expression of BDNF in pyramidal neurons in the PFC and on the restricted expression of α5-GABA_A_R in corticolimbic brain regions, we primarily focused our analysis on the PFC-related molecular and behavioral deficits in the current study. Additional cell type- and brain region-specific approaches will be necessary to further determine the precise role of these molecules. Finally, these studies were performed in male mice and comparative analyses in female mice are warranted.

Nonetheless, given the critical role of p62 in regulating α5-GABA_A_R trafficking and behavioral outcomes in *Bdnf*^+/-^ mice, we propose that these molecular players (*i.e*., p62, BDNF, α5-GABA_A_R) critically contribute to cognitive processes and other brain functions under both normal and pathophysiological conditions. For instance, p62 protein levels typically increase with age, reflecting a gradual decrease in cellular autophagic activity (Vilchez et al., 2014), and elevated p62 levels or increase in p62^+^ inclusions are cardinal features of neurodegenerative disorders, including Alzheimer’s and Parkinson’s diseases (Ferrer et al., 2011; Salminen et al., 2012). Furthermore, we recently reported upregulated expression of p62 proteins in cultured neurons isolated from subjects with schizophrenia and bipolar disorder (Sumitomo et al., 2018a), and in brains of a mouse model of schizophrenia (Sumitomo et al. 2018b). Hence, controlling the dosage of p62 protein may provide a potential target for therapeutic intervention against symptoms shared across these disorders, such as cognitive impairment, through augmentation of inhibitory neurotransmission. Of note, cognitive impairment is among symptoms most difficult to treat and is known to frequently persist during remission in MDD (Disner et al., 2011). The gene expression profiles in MDD in our present study therefore endorse the idea that attenuated autophagy throughout episode/remission states may provide a biological underpinning for persistent cognitive impairment in MDD. In addition, surface availability of GABA_A_ receptors represents a rate-limiting step for GABAergic neurotransmission. Therefore, the mechanisms for regulating surface presentation of GABA_A_ receptor through p62 dosage control may provide an alternative therapeutic approach, for instance for MDD subjects who do not respond to current antidepressant treatment, or for targeting cognitive deficits during remission of depression, across brain disorders and during aging.

## MATERIALS AND METHODS

### Animals

*Bdnf* knockout mice (*Bdnf*^tm1Jae^/J, stock No. 002266) were obtained from the Jackson Laboratory (Bar Harbor, USA). GFP-LC3 transgenic mice were provided by Dr. Noboru Mizushima (University of Tokyo, Japan). p62 knockout mice were provided by Dr. Toru Yanagawa (University of Tsukuba, Japan). CaMKII-p62 transgenic mice were generated according to the standard procedures (Nagy et al., 2003). Mice were maintained on the C57BL/6J genetic background for at least 10 generations. Eight to 12-week old male mice were used for behavioral analysis. Behavioral experiments and data collection were performed by experimenters blind to animal genotypes. Maintenance of mouse colonies and experiments using mice were in accordance with the NIH Guide for the Care and Use of Laboratory Animals, and approved by the Institutional Animal Care and Use Committee at Kyoto University and at the Centre for Addiction and Mental Health.

### Prepulse inhibition

The startle response and prepulse inhibition (PPI) were measured using a startle reflex measurement system (SR-LAB) as described (Sumitomo et al., 2018a). The test session began by placing a mouse in a plastic cylinder and leaving it undisturbed for 30 min. The background white noise level in the chamber was 70 dB. A prepulse–pulse trial started with a 50-ms null period, followed by a 20-ms prepulse white noise (74, 78, 82, 86, or 90 dB). After a 100-ms delay, the startle stimulus (a 40-ms, 120 dB white noise) was presented, followed by a 290-ms recording time. The total duration of each trial was 500 ms. A test session consisted of six trial types (pulse-only trial, and five types of prepulse–pulse trial). Six blocks of the six trial types were presented in a pseudo–randomized order such that each trial type was presented once within a block. The formula: 100– ([Response on acoustic prepulse-pulse stimulus trials/Startle response on pulse-only trials] x 100) was used to calculate %PPI.

### Rule shift assay

Cognitive flexibility was evaluated in the rule shift assay, essentially as described (Bissonette et al., 2008; Cho et al., 2015). In brief, mice were habituated to food, feeding apparatus, different odor cues and digging medium texture cues prior to testing, and then food-deprived a day before the assays. Mice were initially trained in a sequence of trials to associate a food reward with a specific stimulus (*i.e*., either an odor or a digging medium). A varying combination of stimulus and food reward was presented to mice per trial. Eight consecutive correct responses to the food reward were considered reaching criterion (*i.e*., successful establishment of association between the stimulus and the food reward), and the number of trials to reach criterion were scored for each mouse before and after rule shifting (*e.g*., from an odor cue to a different texture cue predicting reward). Upon rule shifting, numbers of errors due to perseveration to an old rule were scored before reaching new criterion.

### Quantitative reverse transcription-polymerase chain reaction (qRT-PCR)

Total RNA was extracted from the PFC using RNeasy Mini kit (Qiagen), and reverse-transcribed with a ReverTra Ace cDNA synthesis kit (Toyobo). TaqMan probes were purchased from Applied Biosystems, Inc. All data were normalized with *Gapdh* as reference.

### Primary cortical neuron culture

Primary cortical neurons were prepared from E13.5 frontal cortex through papain treatment (0.5 μg/ml in Earle’s balanced salt solution supplemented with 5 mM EDTA and 200 μM L-cysteine), followed by mechanical trituration using fire-bore glass pipettes, and plated on poly-_D_-lysine-coated glass cover slips or glass-bottom dishes (MatTek). The cultures were recovered in serum-containing media (Neurobasal media supplemented with 10% horse serum, 5% fetal bovine serum, and 2 mM glutamine [Gibco]) for 4 h and maintained in serum-free media (Neurobasal media supplemented with B-27 (1:50 diluted), 2 mM glutamine, 50 I.U./ml penicillin, and 50 μg/ml streptomycin), with half of media being replaced with fresh media every 2–3 days. The cultures were used for immunofluorescence analysis or for surface biotinylation followed by Western blot analysis.

### Neuronal autophagy and autophagy flux assays

For evaluation of autophagy, primary cortical neurons cultured for 18–25 days were transfected with GFP-LC3 expression plasmid using Lipofectamine 2000 (Invitrogen) via the standard procedure, and 48 h post-transfection, the cells were incubated with BDNF (Sigma, 100 ng/ml) for up to 40 min before fixation with 4% paraformaldehyde (PFA) in PBS. When appropriate, LysoTracker (Invitrogen) was included in culture media 5 min before fixation. Fluorescence images were acquired by confocal microscopy (SP8, Leica) and the fluorescence intensities of GFP+ puncta and LysoTracker uptake were evaluated by ImageJ (NIH). As for the autophagy flux assay (Mizushima et al., 2010), primary cortical neurons were treated with BDNF (100 ng/ml) in the presence or absence of a lysosomal protease inhibitor (bafilomycin A1 [BafA1], 5 ng/ml) and the cell lysates were analyzed by Western blot using LC3 antibody. The amounts of autophagosomal membrane-bound LC3 (LC3-II) normalized by total amounts of LC3 (cytosolic LC3-I plus membrane bound LC3-II) were compared in the presence versus the absence of BafA1 to calculate the autophagy flux, which corresponds to the amount of LC3 degraded through an autophagy-dependent process within the lysosome.

### Surface biotinylation

Biotinylation of cell surface proteins was performed in primary neuron cultures using the cell surface protein isolation kit (Pierce) according to the manufacturer’s protocol. Briefly, cells were incubated with ice-cold PBS containing Sulfo-NHS-SS-Biotin (Pierce) for 30 min with gentle rocking at 4°C. Cells were then lysed and precipitated with NeutrAvidin beads. Precipitated proteins were eluted from the NeutrAvidin beads with loading buffer containing dithiothreitol (DTT) and heated for 5 min at 95°C and then analyzed by Western blot.

### Chemical cross-linking assay

Cell surface receptor cross-linking assays were performed as described (Boudreau et al., 2012). In brief, mice were decapitated and coronal brain slices (~1 mm thick) were quickly prepared using Brain Matrix (Ted Pella) within 30 sec on ice. The PFC was then excised from the slices, minced into small pieces using a razor blade, and incubated in artificial CSF buffer containing bis(sulfosuccinimidyl)suberate (BS3) cross-linker (2 mM, ThermoFisher) for 30 min to 4 h at 4°C with constant invert mixing. After quenching the crosslinking reaction by adding glycine (100 mM) for 10 min at 4°C, the tissues were harvested by centrifugation (20,000 g, 4°C, 2 min). The proteins were prepared in lysis buffer containing 0.1 % Nonidet P-40 (v/v), protease and phosphatase inhibitor cocktail, and 1 mM DTT, and analyzed by Western blot.

### Western blots

Western blot analysis was performed according to the standard procedure. The primary antibodies used were anti-α1-GABA_A_ receptor (rabbit, 1:5,000, abcam), anti-α2-GABA_A_ receptor (rabbit, 1:400, Alomone Labs), anti-α5-GABA_A_ receptor (rabbit, 1:1,000, R&D Systems), anti-NR1 (rabbit monoclonal [1.17.2.6], 1:1,000, Millipore), anti-p62 (guinea pig, 1:1,000, MBL), anti-LC3B (rabbit, 1:1,000, Novus), anti-GAPDH (mouse, 1:1,000, abcam), and anti-α-Tubulin (mouse monoclonal [B-5-1-2], 1:8,000, Sigma).

### Immunofluorescence

Brains of mice (*n* = 4 per group) perfused with 4% PFA/PBS were serially cut into 50 μm-thick coronal sections using vibratome (VT1200S, Leica), and the sections from one cohort of mice (a littermate pair of wild-type and *Bdnf*^+/-^ mice) were permeabilized in PBS containing 0.05% Triton X-100 for 1 h, incubated for 30 min at room temperature in 10% goat serum (Chemicon) in PBS, and immunostained for 16 h at 4°C with primary antibodies followed by Alexa Fluor® 488- or 546-conjugated secondary antibodies (Molecular Probes) for 1 h at room temperature. The stained samples were observed using a confocal microscope (SP8, Leica; 40x objective lens, NA = 1.3); images of one optical section (1 μm thick) were acquired from 3 to 6 non-overlapping areas per section, randomly chosen in PFC (layer 2/3), and 3 to 4 serial sections were analyzed. The fluorescence intensities of p62 immunostaining were measured from each neuronal soma (30~50 somas per section) using ImageJ (NIH), with the background levels of staining in adjacent regions being subtracted, and the average immunofluorescence intensity was calculated across all serial sections from every mouse used. The primary antibodies used were: anti-p62 (guinea pig, 1:400, MBL) and anti-CaMKIIα (mouse monoclonal [6G9], 1:500, Stressmarq).

### Human transcriptome analysis

For differential expression summary statistics, data from a prior meta-analysis of altered gene expression in MDD were used (Ding et al., 2015). In brief, human postmortem brain samples were obtained after consent from next of kin during autopsies conducted at the Allegheny County Medical Examiner’s Office (Pittsburg, PA, USA) using procedures approved by the Institutional Review Board and Committee for Oversight of Research Involving the Dead at the University of Pittsburgh. A total of 51 MDD and 50 control subjects were included in the 8 studies in that report. Samples from the dorsolateral prefrontal cortex (dlPFC), subgenual anterior cingulate cortex (sgACC) or rostral amygdala enriched in lateral, basolateral and basomedial nuclei had been previously collected and processed on Affymetrix HG-U133 Plus 2 or Illumina HT12 gene arrays. Four studies were performed in the sgACC, 2 studies in the amygdala and 2 in the dlPFC. Half of the studies had been performed in female subjects in each brain region. See details on subjects, areas investigated and other parameters in Ding et al., 2015. Differential summary statistics were used to rank the 10,621 genes from the upregulated gene with the lowest p-value to the downregulated gene with lowest p-value. The area under the receiver operating curve statistics was used to test enrichment of the autophagy-related gene lists. Significance between the area under the curve (AUC) values for the autophagy-enhancing and autophagy-attenuating gene lists was empirically determined using 10,000 comparisons of randomly selected gene sets of the same sizes.

For gene set enrichment analysis (GSEA) (Subramanian et al., 2005), RNAseq data on sgACC postmortem samples from four MDD cohorts (single episode, *n* = 20; first remission, *n* = 15; recurrent episode, *n* = 20; and second remission, *n* = 15) and one control cohort (*n* = 20) were used; sample collection procedures, the site of collection and approval body are essentially the same as above (see details in Scifo et al., 2018), and RNAseq procedure and bioinformatics analysis are as described (Shukla et al., 2018). Full datasets on RNAseq analysis will be reported elsewhere (Shukla et al., BioRxiv).

### Statistical analysis

All data were represented as mean ± standard error of the mean (SEM) and were analyzed by Kruskal-Wallis test followed by Dunn’s multiple comparison test, unless otherwise noted. Behavioral assay data were analyzed by one-way or two-way analysis of variance (ANOVA) followed by Bonferroni post-hoc test, using Prism statistics software (GraphPad).

## ACKNOWLEDGMENTS

This work was supported by grants from the Canadian Institute of Health Research (CIHR #153175 to E.S.), National Alliance for Research on Schizophrenia and Depression (NARSAD award #25637 to E.S.), the National Institutes of Health (MH-093723 to E.S.), Campbell Family Mental Health Research Institute (to E.S.), and Department of Defense/Congressionally Directed Medical Research Program (W81XWH-11-1-0269 to T.T.).

## AUTHOR CONTRIBUTIONS

T.T. and E.S. conceived the studies; T.T., A.S., and Y.H.-T. carried out experiments; H.M. generated CaMKII-p62 transgenic mice; H.O. provided analytical tools; R.S. and L.F. analyzed gene expression profiles; T.T. and E.S. wrote and edited the paper.

## Supplemental Information

**Figure S1.**
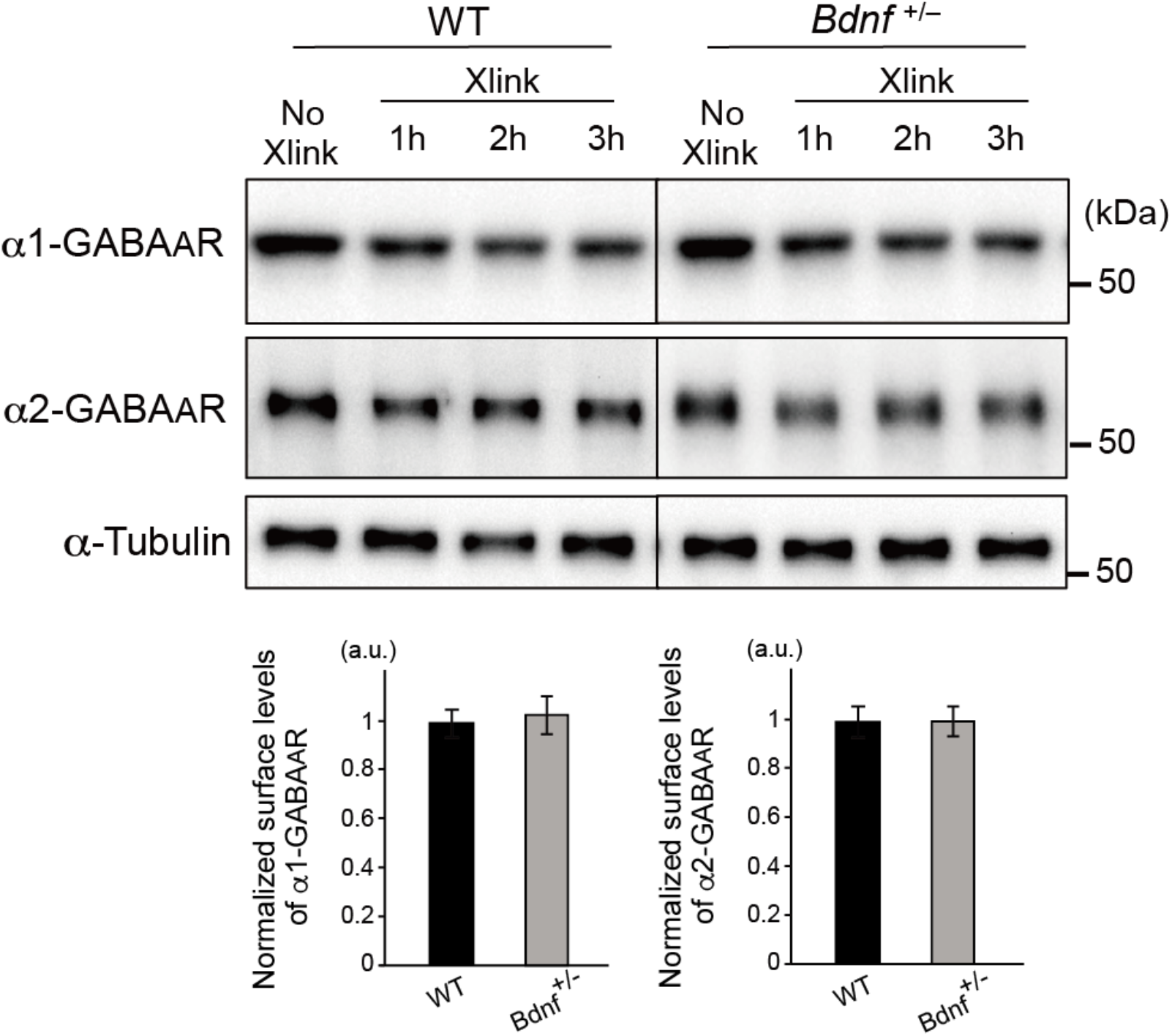
Surface presentation of α1- and α2-GABA_A_R in *Bdnf*^+/-^ mice (Related to Figure 2) BS3 cross-linking assays using the PFC extracts from WT (*n* = 3) and *Bdnf*^+/-^ mice (*n* = 3). Levels of non-crosslinked α1- and α2-GABA_A_R (~50 kDa) were normalized to levels of α-Tubulin at each time point. Differences in the levels of non-crosslinked α1- or α2-GABA_A_R at a given time point versus those of the control sample (No Xlink: no cross-linker added) represent the amounts of surface GABA_A_R that underwent mobility shift toward a higher molecular weight range due to covalent crosslinking with anonymous cell surface proteins. No significant differences in the surface levels of α1- and α2-GABA_A_R were observed in the PFC of WT versus *Bdnf*^+/-^ mice.

**Figure S2.**
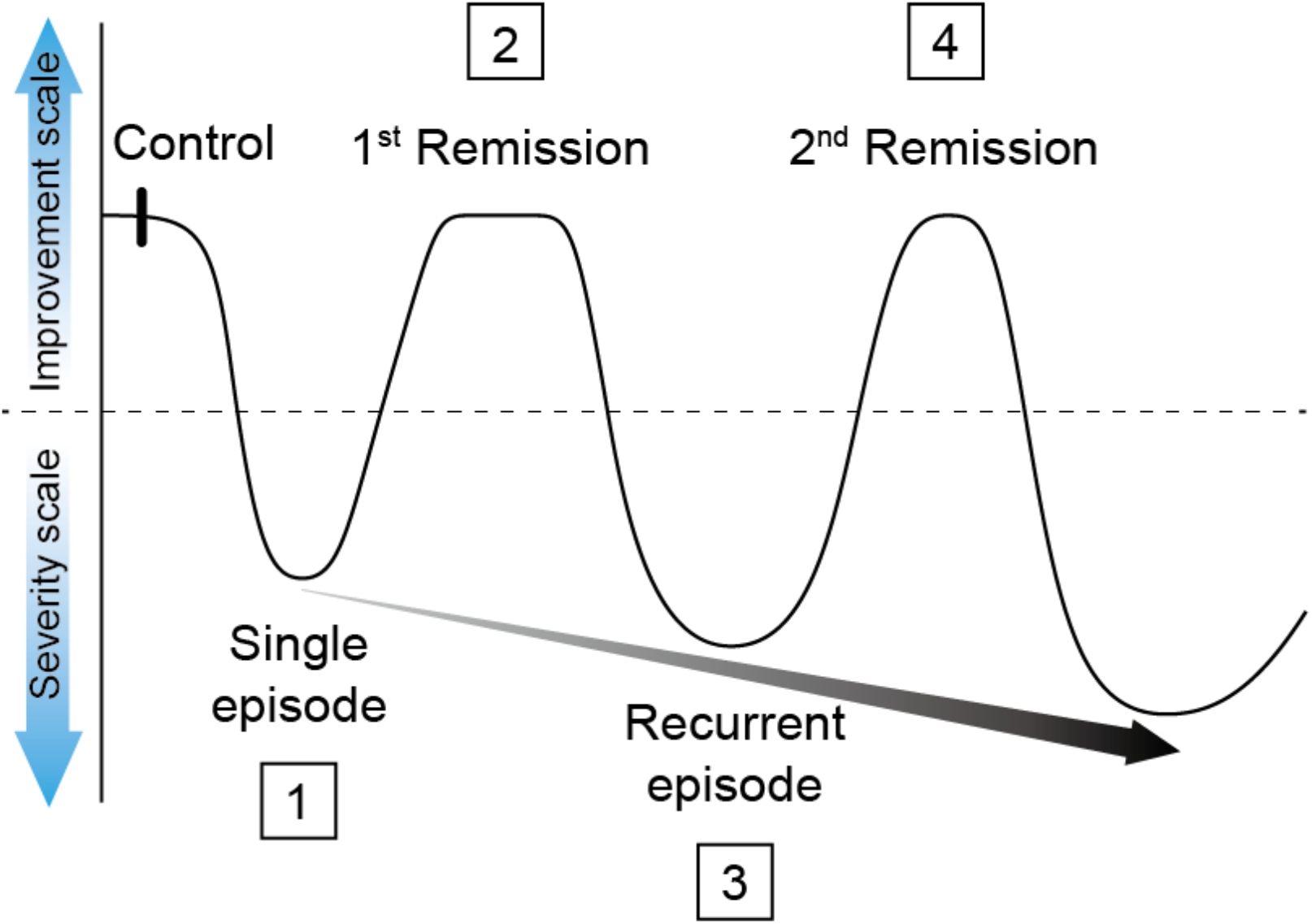
Overview of MDD and control cohorts used in RNAseq study (Related to Figure 4) Predicted trajectory of MDD pathology is shown across various disease states, depicting recurring episodes of increasing severity (gray arrow) and shorter remission periods. 1. MDD single episode; 2. MDD first remission; 3. MDD recurrent episode; and 4. MDD second remission. Reproduced and modified with permission from Sibille and French (2013).

**Figure S3.**
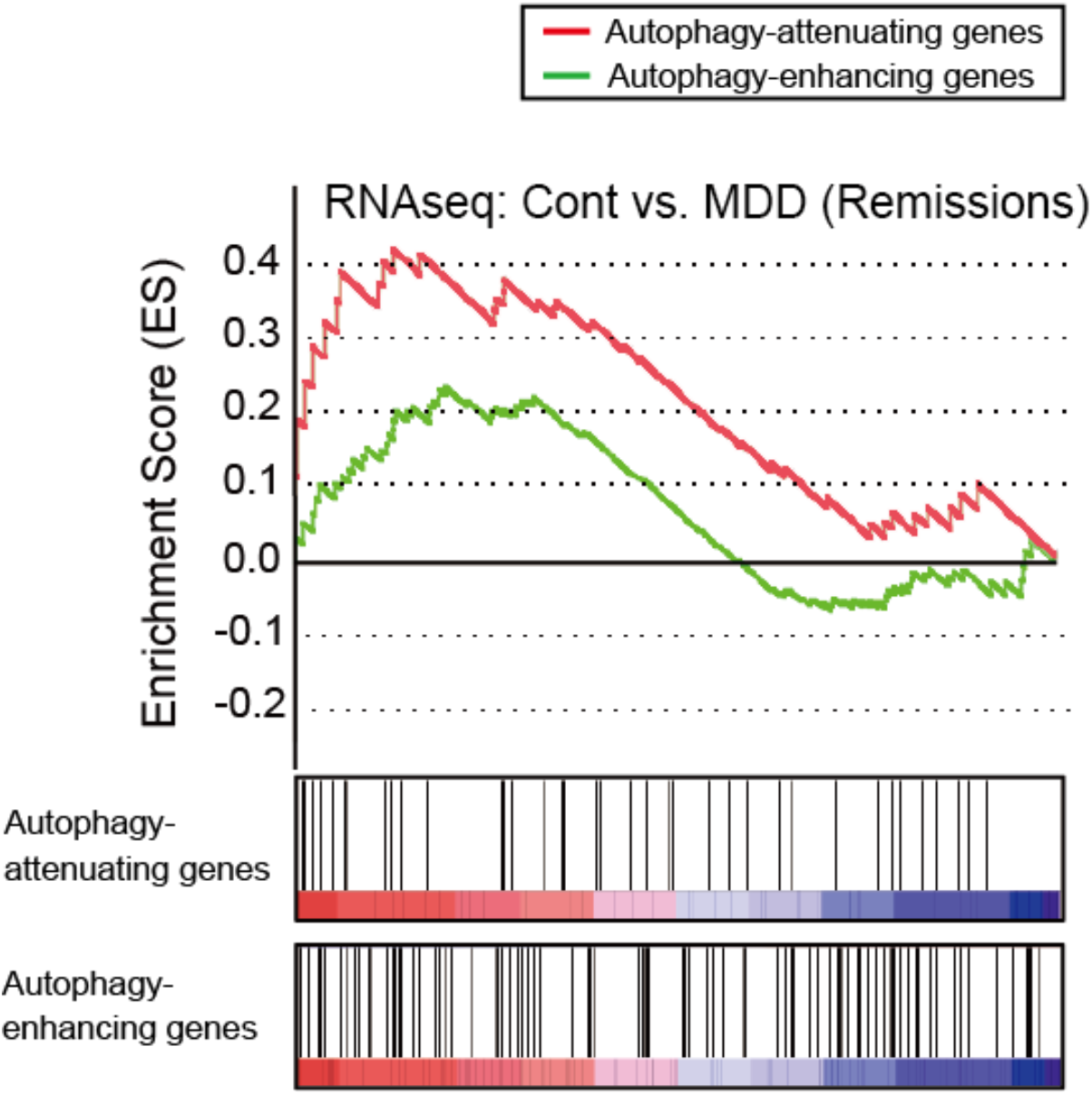
GSEA on autophagy gene expression in MDD remission states (Related to Figure 4) GSEA on gene expression profiles in MDD (1^st^ and 2^nd^ remission states combined) versus control subjects obtained via RNAseq shows a significant enrichment in upregulation for autophagy-attenuating genes (red line) (p=0.016) but no significant enrichment for autophagy-enhancing genes (green line).

**Figure S4.**
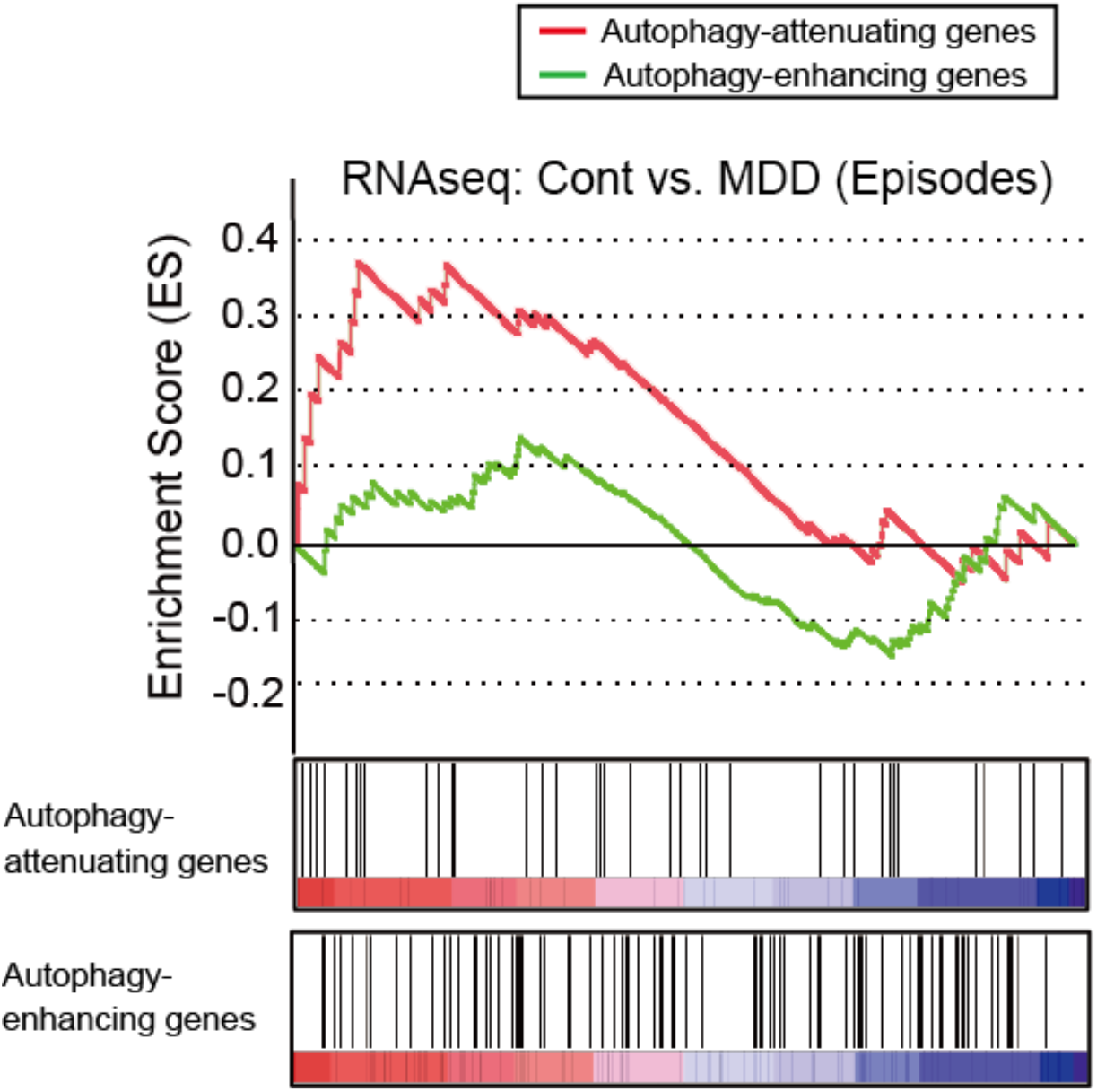
GSEA on autophagy gene expression in MDD episode states (Related to Figure 4) GSEA on gene expression profiles in MDD (single and recurrent episode states combined) versus control subjects obtained via RNAseq shows the trend level of enrichment in upregulation for autophagy-attenuating genes (red line) (p=0.098) but no significant enrichment for autophagy-enhancing genes (green line).

**Table S1.**
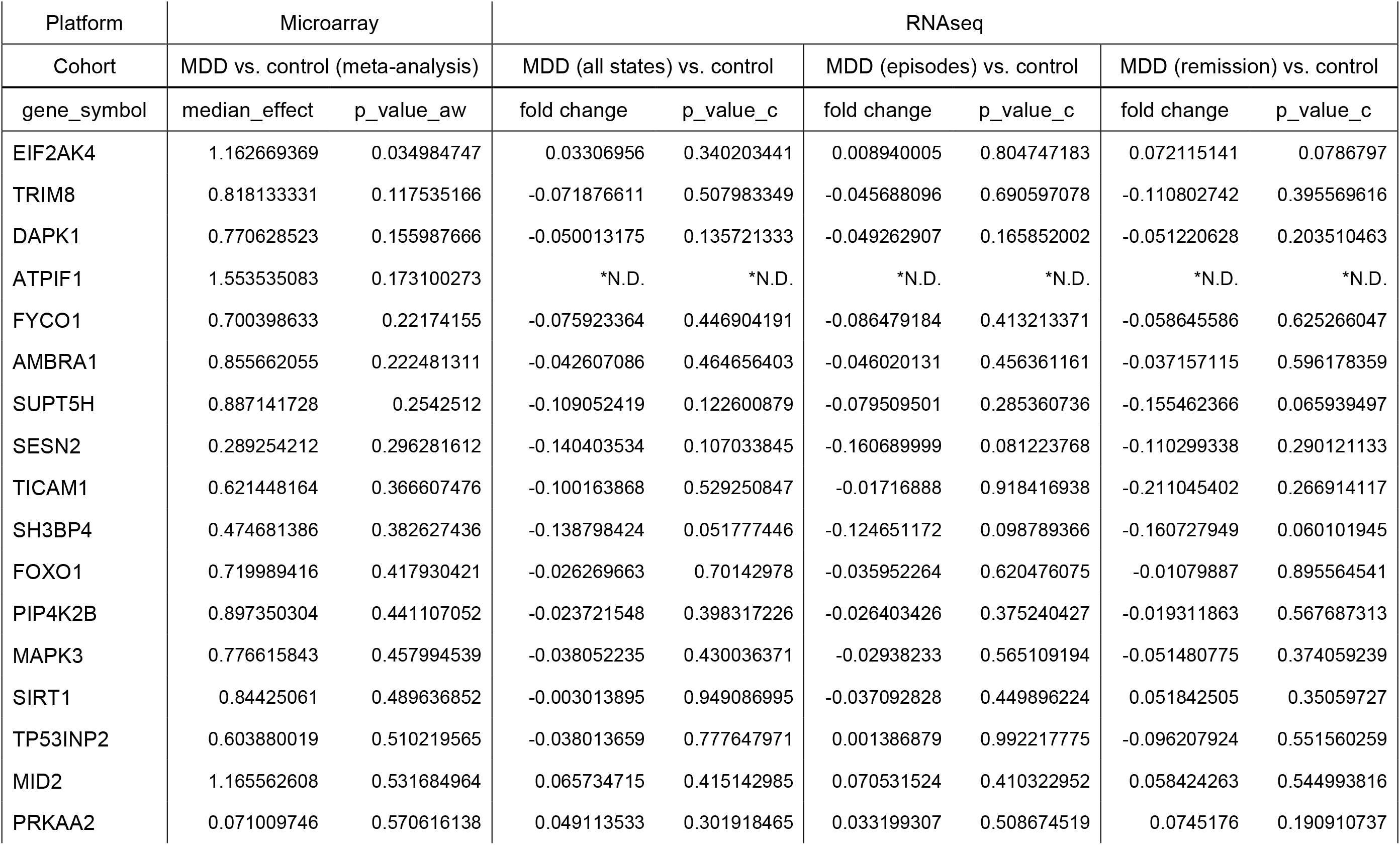
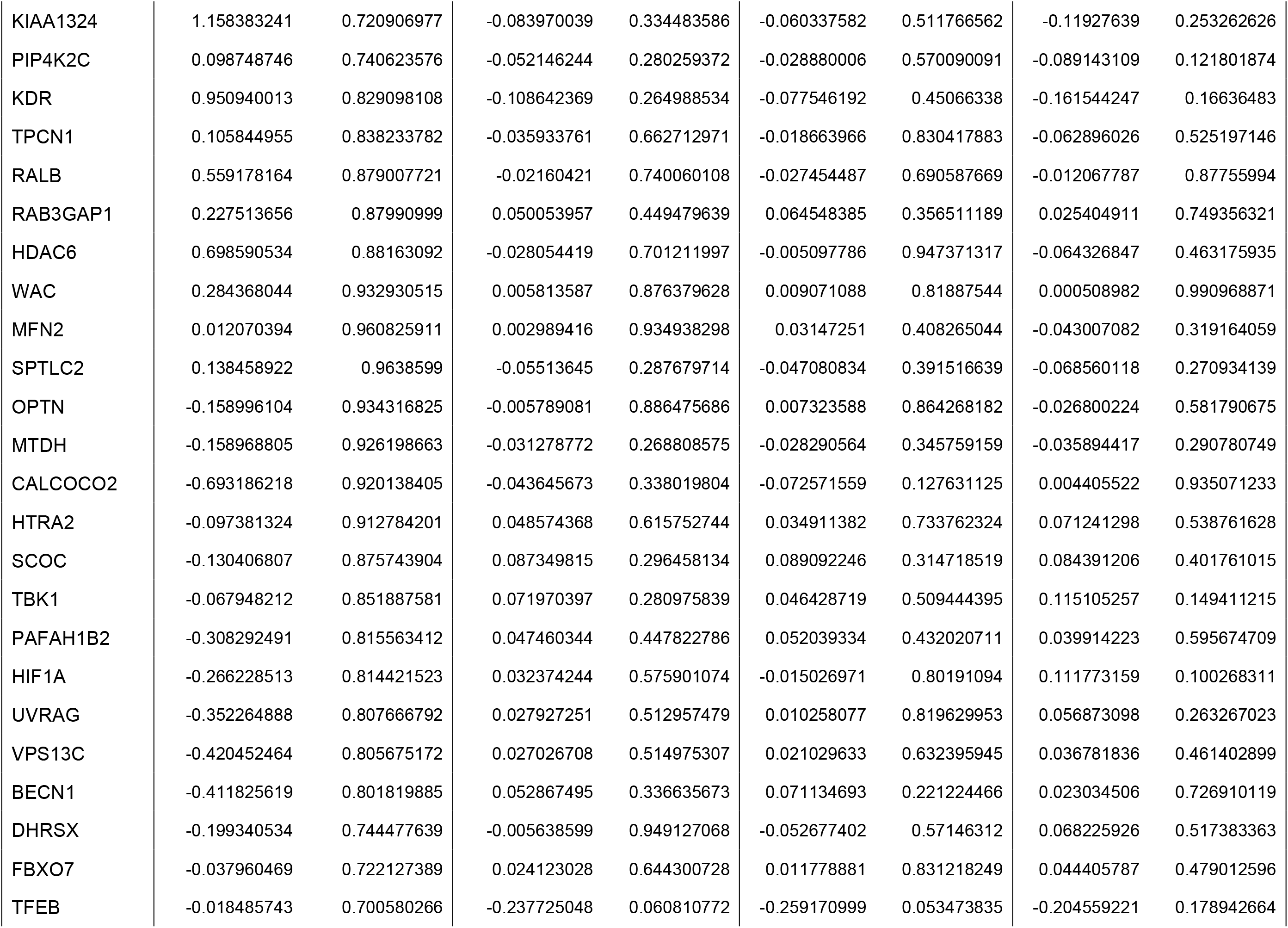
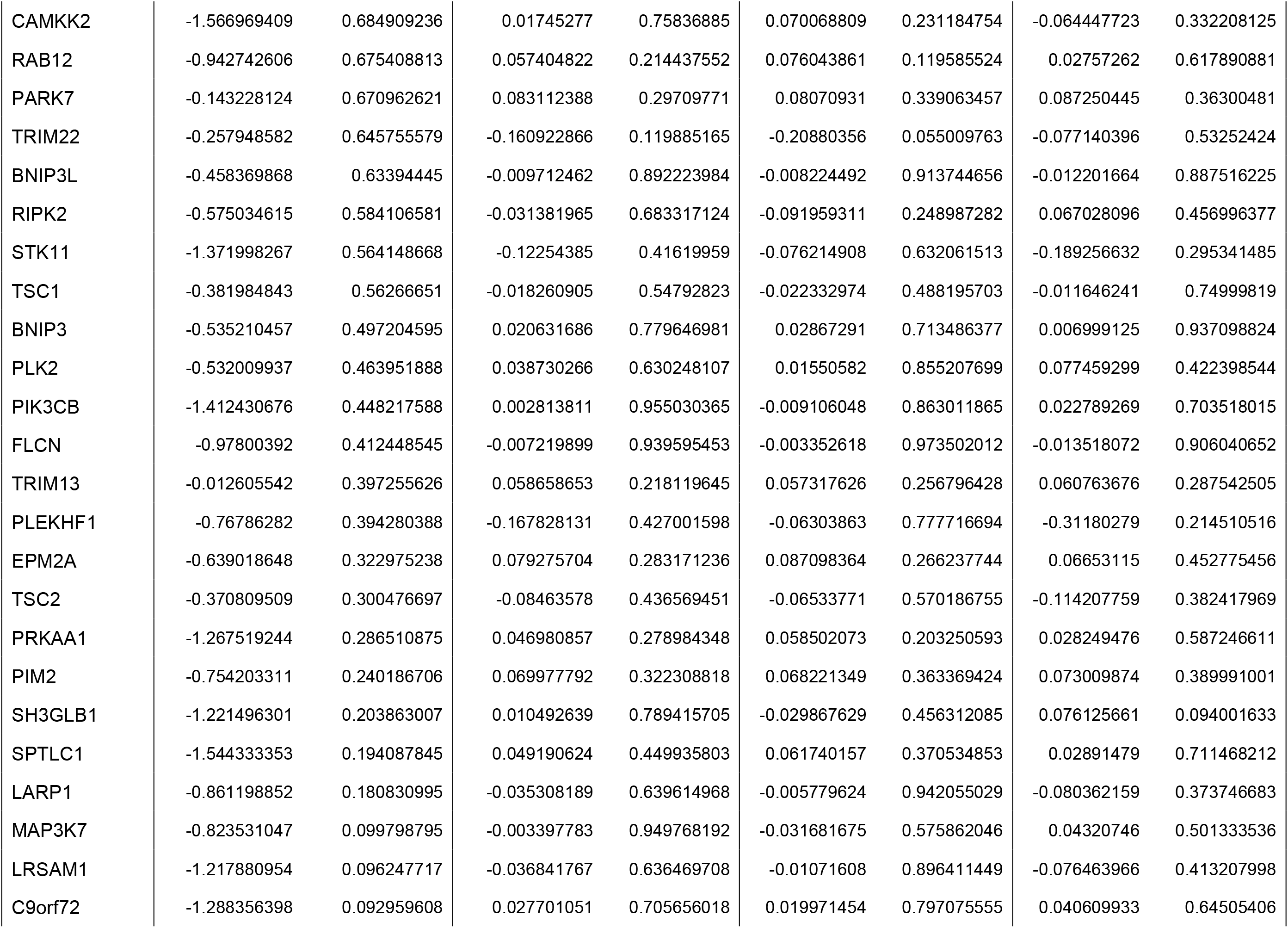
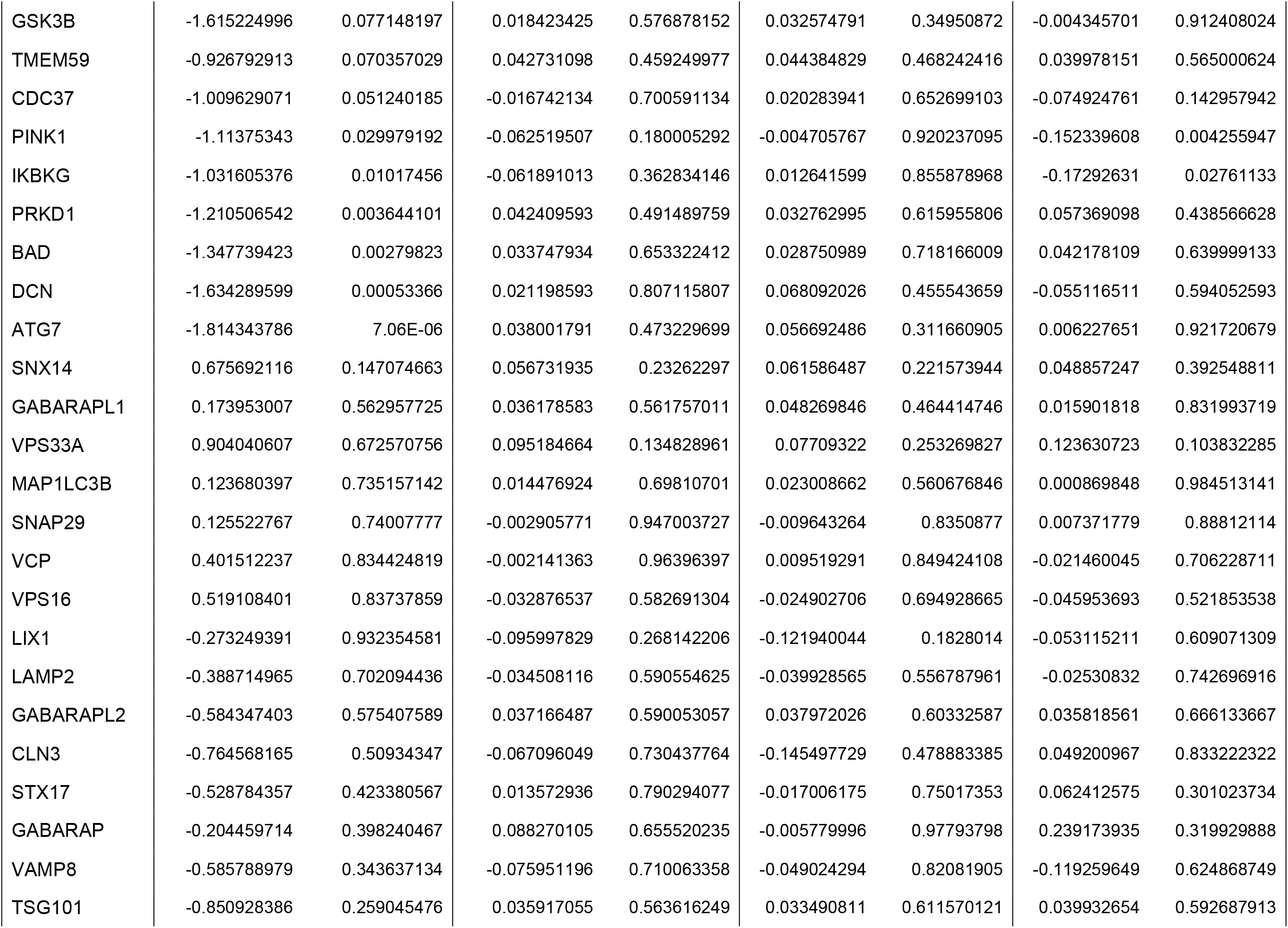
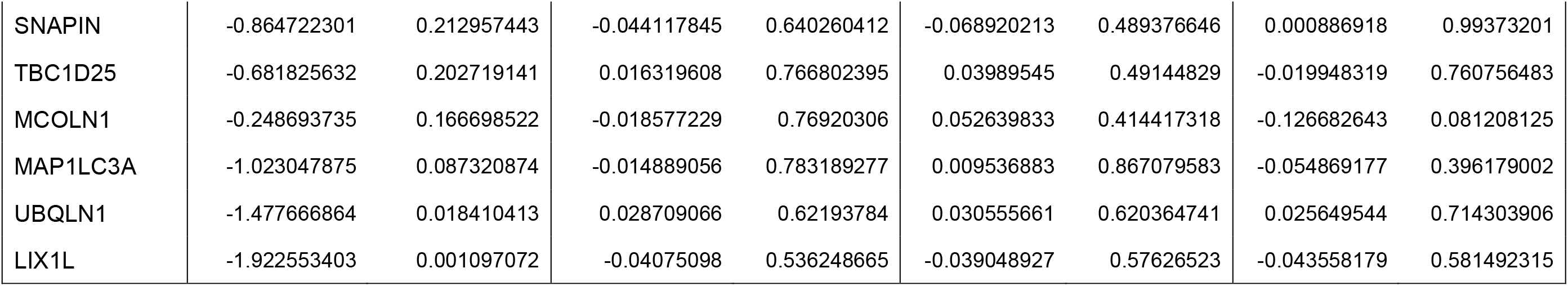
Autophagy-enhancing gene list (Related to Figure 4, Figure S3, Figure S4) Due to overlap of genes among related GO term for autophagy-related genes, GO term “Positive regulation of autophagy” and GO term “Autophagy maturation” were combined to generate the list of “Autophagy-enhancing genes” (95 genes in total). *N.D.: Not detected.

**Table S2.**
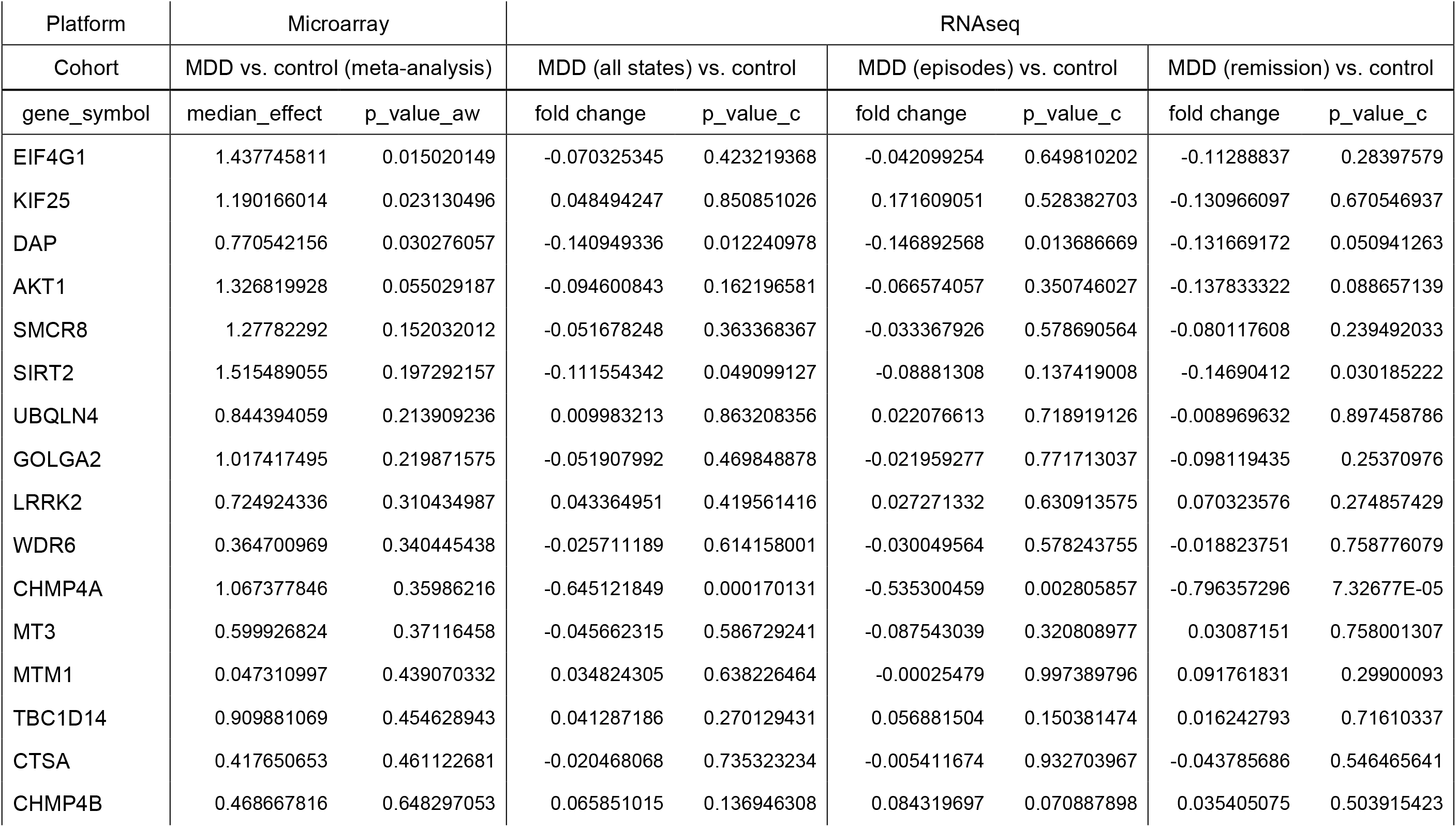
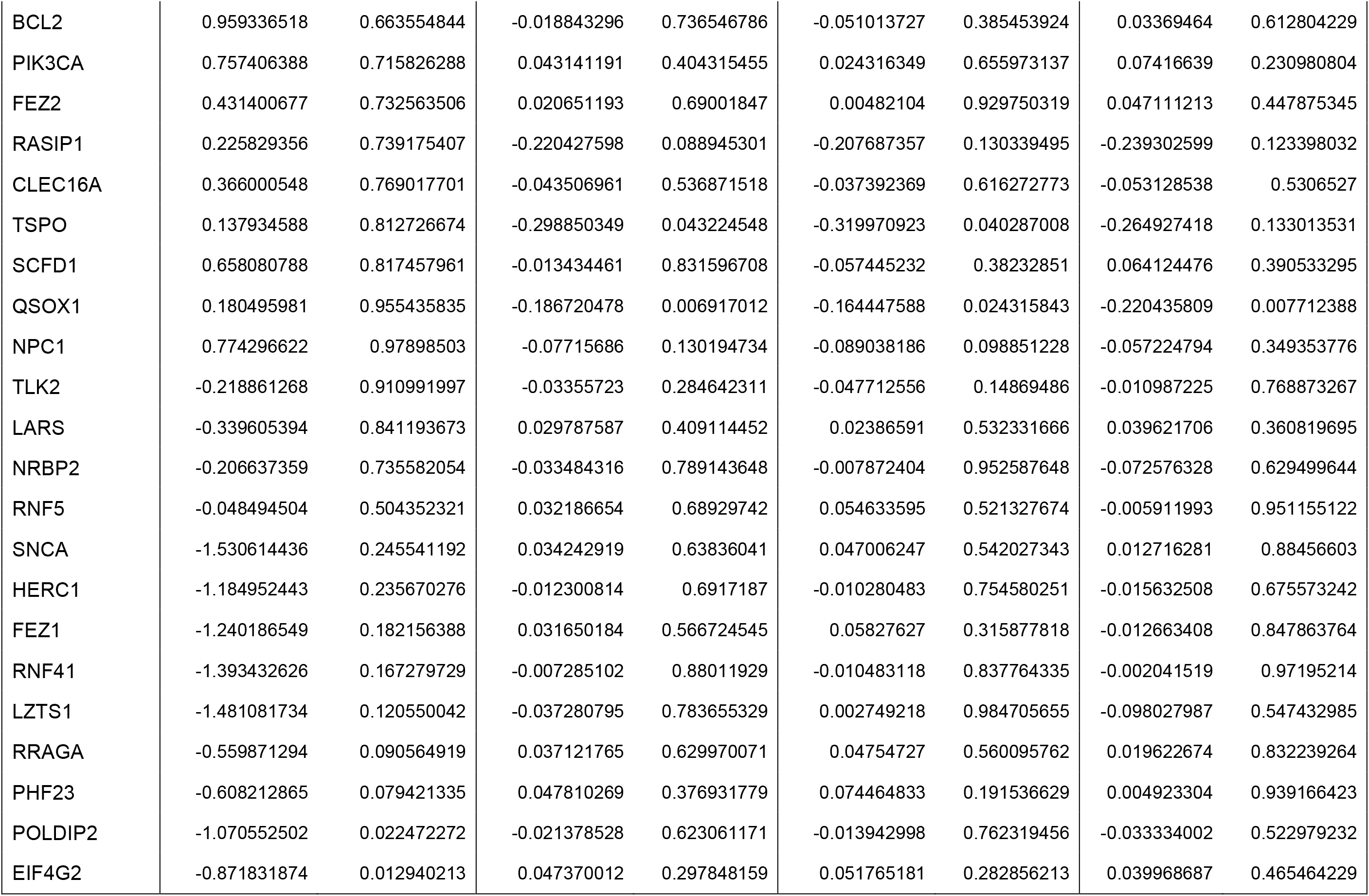
Autophagy-attenuating gene list (Related to Figure 4, Figure S3, Figure S4) Due to overlap of genes between GO term “Positive regulation of autophagy” and GO term “Negative regulation of autophagy”, four genes (*ATG7, TSC1, TSC2, PINK1*) that are classified as positive regulators of autophagy in the vast majority of literature were removed from the list of GO term “Negative regulation of autophagy” to generate the list of “Autophagy-attenuating genes” (38 genes in total).

## REFERENCES

Bast, T., Pezze, M., and McGarrity, S. (2017). Cognitive deficits caused by prefrontal cortical and hippocampal neural disinhibition. Br. J. Pharmacol. 174, 3211–3225.

Bissonette, G.B., Martins, G.J., Franz, T.M., Harper, E.S., Schoenbaum, G., and Powell, E.M. (2008). Double dissociation of the effects of medial and orbital prefrontal cortical lesions on attentional and affective shifts in mice. J. Neurosci. 28, 11124–11130.

Boudreau, A.C., Milovanovic, M., Conrad, K.L., Nelson, C., Ferrario, C.R., and Wolf, M.E. (2012). A protein crosslinking assay for measuring cell surface expression of glutamate receptor subunits in the rodent brain after in vivo treatments. Curr. Protoc. Neurosci. CHAPTER: Unit–5.3019.

Caballero, A., and Tseng, K.Y. (2016). GABAergic function as a limiting factor for prefrontal maturation during adolescence. Trends Neurosci. 39, 441–448.

Calabrese, F., Guidotti, G., Racagni, G., and Riva, M.A. (2013). Reduced neuroplasticity in aged rats: a role for the neurotrophin brain-derived neurotrophic factor. Neurobiol. Aging 34, 2768–2776.

Chao, M.V. (2003). Neurotrophins and their receptors: a convergence point for many signalling pathways. Nat. Rev. Neurosci. 4, 299–309.

Chao, M.V., Rajagopal, R., and Lee, F.S. (2006). Neurotrophin signalling in health and disease. Clin. Sci. (Lond) 110, 167–173.

Cho, K.K., Hoch, R., Lee, A.T., Patel, T., Rubenstein, J.L., and Sohal, V.S. (2015). Gamma rhythms link prefrontal interneuron dysfunction with cognitive inflexibility in Dlx5/6^+/-^ mice. Neuron 85, 1332–1343.

Dincheva, I., Glatt, C.E., and Lee, F.S. (2012). Impact of the BDNF Val66Met polymorphism on cognition: implications for behavioral genetics. Neuroscientist 18, 439–451.

Dincheva, I., Lynch, N.B., and Lee, F.S. (2016). The role of BDNF in the development of fear learning. Depress. Anxiety 33, 907–916.

Ding, Y., Chang, L.C., Wang, X., Guilloux, J.P., Parrish, J., Oh, H., French, B.J., Lewis, D.A., Tseng, G.C., and Sibille, E. (2015). Molecular and genetic characterization of depression: overlap with other psychiatric disorders and aging. Mol. Neuropsychiatry 1, 1–12.

Disner, S.G., Beevers, C.G., Haigh, E.A., and Beck, A.T. (2011). Neural mechanisms of the cognitive model of depression. Nat. Rev. Neurosci. 12, 467–477.

Engin, E., Zarnowska, E.D., Sigal, M., Keist, R., Zeller, A., Pearce, R.A., and Rudolph, U. (2013). Alpha5-containing GABAA receptors in dentate gyrus enable cognitive flexibility. FASEB J. 27, 661.7–661.7.

Fee, C., Banasr, M., and Sibille, E. (2017). Somatostatin-positive gamma-aminobutyric acid interneuron deficits in depression: cortical microcircuit and therapeutic perspectives. Biol. Psychiatry 82, 549–559.

Ferrer, I., Martinez, A., Blanco, R., Dalfó, E., and Carmona, M. (2011). Neuropathology of sporadic Parkinson disease before the appearance of parkinsonism: preclinical Parkinson disease. J. Neural Transm. (Vienna) 118, 821–839.

Fritschy, J.M., and Mohler, H. (1995). GABAA-receptor heterogeneity in the adult rat brain: differential regional and cellular distribution of seven major subunits. J. Comp. Neurol. 1359, 154–194.

Gorba, T., and Wahle, P. (1999). Expression of TrkB and TrkC but not BDNF mRNA in neurochemically identified interneurons in rat visual cortex *in vivo* and in organotypic cultures. Eur. J. Neurosci. 11, 1179–1190.

Guilloux, J.P., Douillard-Guilloux, G., Kota, R., Wang, X., Gardier, A.M., Martinowich, K., Tseng, G.C., Lewis, D.A., and Sibille, E. (2012). Molecular evidence for BDNF- and GABA-related dysfunctions in the amygdala of female subjects with major depression. Mol. Psychiatry 17, 1130–1142.

Hauser, J., Rudolph, U., Keist, R., Möhler, H., Feldon, J., and Yee, B.K. (2005). Hippocampal alpha5 subunit-containing GABAA receptors modulate the expression of prepulse inhibition. Mol. Psychiatry 10, 201–207.

Hoftman, G.D., Datta, D., and Lewis, D.A. (2017). Layer 3 Excitatory and inhibitory circuitry in the prefrontal cortex: developmental trajectories and alterations in schizophrenia. Biol. Psychiatry 81, 862–873.

Huang, D.W., Sherman, B.T., and Lempicki, R.A. (2009). Systematic and integrative analysis of large gene lists using DAVID Bioinformatics Resources. Nat. Protoc. 4, 44–57.

Islam, F., Mulsant, B.H., Voineskos, A.N., and Rajji, T.K. (2017). Brain-derived neurotrophic factor expression in individuals with schizophrenia and healthy aging: testing the accelerated aging hypothesis of schizophrenia. Curr. Psychiatry Rep. 19, 36.

Kellendonk, C., Simpson, E.H., and Kandel, E.R. (2009). Modeling cognitive endophenotypes of schizophrenia in mice. Trends Neurosci. 32, 347–358.

Knight, M.J., and Baune, B.T. (2018). Cognitive dysfunction in major depressive disorder. Curr. Opin. Psychiatry 31, 26–31.

Kononenko, N.L., Claßen, G.A., Kuijpers, M., Puchkov, D., Maritzen, T., Tempes, A., Malik, A.R., Skalecka, A., Bera, S., Jaworski, J., and Haucke, V. Retrograde transport of TrkB-containing autophagosomes via the adaptor AP-2 mediates neuronal complexity and prevents neurodegeneration. Nat. Commun. 8, 14819.

Laudes, T., Meis, S., Munsch, T., and Lessmann, V. (2012). Impaired transmission at corticothalamic excitatory inputs and intrathalamic GABAergic synapses in the ventrobasal thalamus of heterozygous BDNF knockout mice. Neuroscience 222, 215–227.

Lewis, D.A., Hashimoto, T., and Volk, D.W. (2005). Cortical inhibitory neurons and schizophrenia. Nat. Rev. Neurosci. 6, 312–324.

Lippai, M., and Lőw, P. (2014). The role of the selective adaptor p62 and ubiquitin-like proteins in autophagy. Biomed. Res. Int. 2014, 832704.

Lu, B., Nagappan, G., and Lu, Y. (2014). BDNF and synaptic plasticity, cognitive function, and dysfunction. Handb. Exp. Pharmacol. 220, 223–250.

Manning, E.E., and van den Buuse, M. (2013). BDNF deficiency and young-adult methamphetamine induce sex-specific effects on prepulse inhibition regulation. Front. Cell. Neurosci. 7, 92.

Mizushima, N., Yamamoto, A., Matsui, M., Yoshimori, T., and Ohsumi, Y. (2004). In vivo analysis of autophagy in response to nutrient starvation using transgenic mice expressing a fluorescent autophagosome marker. Mol. Biol. Cell 15, 1101–1111.

Mizushima, N., Yoshimori, T., and Levine, B. (2010). Methods in mammalian autophagy research. Cell 140, 313–326.

Mizushima, N., and Komatsu, M. (2011). Autophagy: renovation of cells and tissues. Cell 147, 728–741.

Nagy, A., Gertsenstein, M., Vintersten, K., and Behringer, R. (2003). Manipulating the mouse embryo. A Laboratory Manual. 3^rd^ Edition. Cold Spring Harbor Laboratory Press.

Negrón-Oyarzo, I., Aboitiz, F., and Fuentealba, P. (2016). Impaired functional connectivity in the prefrontal cortex: a mechanism for chronic stress-induced neuropsychiatric disorders. Neural Plast. 2016, 7539065.

Northoff, G., and Sibille, E. (2014). Cortical GABA neurons and self-focus in depression: a model linking cellular, biochemical and neural network findings. Mol. Psychiatry 19, 959.

Oh, H., Lewis, D.A., and Sibille, E. (2016). The role of BDNF in age-dependent changes of excitatory and inhibitory synaptic markers in the human prefrontal cortex. Neuropsychopharmacology 41, 3080–3091.

Oh, H., Piantadosi, S.C., Rocco, B.R., Lewis, D.A., Watkins, S.C., and Sibille, E. (2018). The role of dendritic brain-derived neurotrophic factor transcripts on altered inhibitory circuitry in depression. Biol. Psychiatry doi: 10.1016/j.biopsych.2018.09.026.

Pankiv, S., Clausen, T.H., Lamark, T., Brech, A., Bruun, J.A., Outzen, H., Øvervatn, A., Bjørkøy, G., and Johansen, T. (2007). p62/SQSTM1 binds directly to Atg8/LC3 to facilitate degradation of ubiquitinated protein aggregates by autophagy. J. Biol. Chem. 282, 24131–24145.

Parikh, V., Naughton, S.X., Yegla, B., and Guzman, D.M. (2016). Impact of partial dopamine depletion on cognitive flexibility in BDNF heterozygous mice. Psychopharmacol. (Berl) 233, 1361–1375.

Parnaudeau, S., O’Neill, P.K., Bolkan, S.S., Ward, R.D., Abbas, A.I., Roth, B.L., Balsam, P.D., Gordon, J.A., and Kellendonk, C. (2013). Inhibition of mediodorsal thalamus disrupts thalamofrontal connectivity and cognition. Neuron 77, 1151–1162.

Pillai, A., Kale, A., Joshi, S., Naphade, N., Raju, M.S., Nasrallah, H., and Mahadik, S.P. (2010). Decreased BDNF levels in CSF of drug-naive first-episode psychotic subjects: correlation with plasma BDNF and psychopathology. Int. J. Neuropsychopharmacol. 13, 535–539.

Porges, E.C., Woods, A.J., Edden, R.A., Puts, N.A., Harris, A.D., Chen, H., Garcia, A.M., Seider, T.R., Lamb, D.G., Williamson, J.B., and Cohen, R.A. (2017). Frontal gamma-aminobutyric acid concentrations are associated with cognitive performance in older adults. Biol. Psychiatry Cogn. Neurosci. Neuroimaging 2, 38–44.

Salminen, A., Kaarniranta, K., Haapasalo, A., Hiltunen, M., Soininen, H., and Alafuzoff, I. (2012). Emerging role of p62/sequestosome-1 in the pathogenesis of Alzheimer’s disease. Prog. Neurobiol. 96, 87–95.

Sánchez-Sánchez, J., and Arévalo, J.C. (2017). A review on ubiquitination of neurotrophin receptors: facts and perspectives. Int. J. Mol. Sci. 18, pii: E630.

Scifo, E., Pabba, M., Kapadia, F., Ma, T., Lewis, D.A., Tseng, G.C., and Sibille, E. (2018). Sustained molecular pathology across episodes and remission in major depressive disorder. Biol. Psychiatry 83, 81–89.

Shukla, R., Prevot, T.D., French, L., Isserlin, R., Rocco, B.R., Banasr, M., Bader, G.D., and Sibille, E. (2018). The relative contributions of cell-dependent cortical microcircuit aging to cognition and anxiety. Biol. Psychiatry doi: 10.1016/j.biopsych.2018.09.019.

Sibille, E., and French, B. (2013). Biological substrates underpinning diagnosis of major depression. Int J Neuropsychopharmacol. 16, 1893–1909.

Subramanian, A., Tamayo, P., Mootha, V.K., Mukherjee, S., Ebert, B.L., Gillette, M.A., et al. (2005). Gene set enrichment analysis: a knowledge-based approach for interpreting genome-wide expression profiles. Proc. Natl. Acad. Sci. U.S.A. 102, 15545–15550.

Sumitomo, A., Yukitake, H., Hirai, K., Horike, K., Ueta, K., Chung, Y., Warabi, E., Yanagawa, T, Kitaoka, S., Furuyashiki, T., Narumiya, S., Hirano, T., Niwa, M., Sibille, E., Hikida, T., Sakurai, T., Ishizuka, K., Sawa, A., and Tomoda, T. (2018a). Ulk2 controls cortical excitatory-inhibitory balance via autophagic regulation of p62 and GABAA receptor trafficking in pyramidal neurons. Hum. Mol. Genet. 27, 3165–3176.

Sumitomo, A., Horike, K., Hirai, K., Butcher, N., Boot, E., Sakurai, T., Nucifora, F.C. Jr., Bassett, A.S., Sawa, A., and Tomoda, T. (2018b). A mouse model of 22q11.2 deletions: Molecular and behavioral signatures of Parkinson’s disease and schizophrenia. Sci. Adv. 4, eaar6637.

Swerdlow, N.R., Braff, D.L., and Geyer, M.A. (2016). Sensorimotor gating of the startle reflex: what we said 25 years ago, what has happened since then, and what comes next. J. Psychopharmacol. 30, 1072–1081.

Tang, G., Gudsnuk, K., Kuo, S.H., Cotrina, M.L., Rosoklija, G., Sosunov, A., Sonders, M.S., Kanter, E., Castagna, C., Yamamoto, A., Yue, Z., Arancio, O., Peterson, B.S., Champagne, F., Dwork, A.J., Goldman, J., and Sulzer, D. (2014). Loss of mTOR-dependent macroautophagy causes autistic-like synaptic pruning deficits. Neuron 83, 1131–1143.

Tripp, A., Oh, H., Guilloux, J.P., Martinowich, K., Lewis, D.A., and Sibille, E. (2012). Brain-derived neurotrophic factor signaling and subgenual anterior cingulate cortex dysfunction in major depressive disorder. Am. J. Psychiatry 169, 1194–1202.

Vilchez, D., Saez, I., and Dillin, A. (2014). The role of protein clearance mechanisms in organismal ageing and age-related diseases. Nat. Commun. 5, 5659.

Wang, H., Bedford, F.K., Brandon, N.J., Moss, S.J., and Olsen, R.W. (1999). GABA(A)-receptor-associated protein links GABA(A) receptors and the cytoskeleton. Nature 397, 69–72.

Weickert, C.S., Hyde, T.M., Lipska, B.K., Herman, M.M., Weinberger, D.R., and Kleinman, J.E. (2003). Reduced brain-derived neurotrophic factor in prefrontal cortex of patients with schizophrenia. Mol. Psychiatry 8, 592–610.

